# A long read optimized *de novo* transcriptome pipeline reveals novel ocular developmentally regulated gene isoforms and disease targets

**DOI:** 10.1101/2020.08.21.261644

**Authors:** Vinay S. Swamy, Temesgen D. Fufa, Robert B. Hufnagel, David M. McGaughey

## Abstract

*De novo* transcriptome construction from short-read RNA-seq is a common method for reconstructing mRNA transcripts within a given sample. However, the precision of this process is unclear as it is difficult to obtain a ground-truth measure of transcript expression. With advances in third generation sequencing, full length transcripts of whole transcriptomes can be accurately sequenced to generate a ground-truth transcriptome. We generated long-read PacBio and short-read Illumina RNA-seq data from a human induced pluripotent stem cell- derived retinal pigmented epithelium (iPSC-RPE) cell line. We use long-read data to identify simple metrics for assessing *de novo* transcriptome construction and optimize a short-read based *de novo* transcriptome construction pipeline. We apply this this pipeline to construct transcriptomes for 340 short-read RNA-seq samples originating from healthy adult and fetal human retina, cornea, and RPE. We identify hundreds of novel gene isoforms and examine their significance in the context of ocular development and disease.

## Introduction

The transcriptome is defined as the set of unique RNA transcripts expressed in a biological system. A single gene can have multiple distinct transcripts, or isoforms, and there are multiple biological processes that drive the formation of these isoforms including alternative promoter usage, alternative splicing, and alternative polyadenylation. Gene isoforms can have distinct and critical functions in biological processes like development, cell differentiation, and cell migration (Dykes et al., 2018), (Trapnell et al., 2010), (Mitra et al., 2020). Alternative usage of isoforms has also been implicated in multiple diseases including cancer, cardiovascular disease, Alzheimer’s disease and diabetic retinopathy (Vitting-Seerup and Sandelin, 2017), (Neagoe Ciprian et al., 2002), (Mills et al., 2013), (Perrin et al., 2005).

Accurate annotation of gene isoforms is fundamental for understanding their biological impact. For example, while the Gencode human comprehensive transcript annotation (release 28) contains 82335 protein coding and 121500 noncoding transcripts across 19901 genes and 38480 pseudogenes, but this annotation is incomplete (Frankish et al., 2019), (Zhang et al., 2020). Some of the first high throughput methods to find novel gene isoforms used short-read (∼100bp) RNA-seq to identify novel exon-exon junctions and novel exon boundaries based soley on RNA-seq coverage (Nagalakshmi et al., 2008). More recently, several groups have developed specialized tools to use RNA-seq to reconstruct the whole transcriptome of a biological sample, dubbed *de novo* transcriptome construction (Haas et al., 2013),(Trapnell et al., 2010), (Pertea et al., 2015).

*De novo* transcriptome construction uses short-read RNA-seq to reconstruct full-length mRNA transcripts. However, a large number of samples are necessary to overcome the noise and short-read lengths of this type of data. Because of increasingly inexpensive sequencing cost, datasets of the necessary size are now available. For example, one of the most comprehensive *de novo* transcriptome projects to date is CHESS, which uses the GTEx data set to construct *de novo* transcriptomes in over 9000 RNA-seq samples from 44 distinct body locations to create a comprehensive annotation of mRNA transcripts across the human body (GTEx Consortium et al., 2017), (Pertea et al., 2018). However, since the GTEx dataset does not include samples from any ocular tissues, the CHESS database remains an incomplete annotation of the human transcriptome.

Despite the increasing number of tools developed, there is no gold standard to evaluate the precision and sensitivity of *de novo* transcriptome construction on real (not simulated) biological data. Long-read sequencing technologies provide a potential solution to this problem as long-read sequencing can capture full length transcripts and thus, can be used to identify a more comprehensive range of gene isoforms. While previous iterations of long-read sequencing technologies typically had higher error rates, the new PacBio Sequel II system sequences long-reads as accurately as short-read based sequencing (Wenger et al., 2019).

We propose that long-read based transcriptomes can serve as a ground truth for evaluating short-read based transcriptomes. In this study, we used PacBio long-read RNA sequencing to inform the construction of short-read transcriptomes. We generated PacBio long-read RNA-seq along with matched Illumina short-read RNA-seq data from a human induced pluripotent stem cell (iPSC)-differentiated retinal pigmented epithelium (RPE) cell line. We then designed a rigorous StringTie-based pipeline that maximizes the concordance between short and long-read *de novo* transcriptomes.

Finally, we applied this optimized pipeline to a data set containing 340 human ocular tissue samples compiled from mining previously published, publicly available short-read RNA-seq data (Swamy and McGaughey, 2019). We built transcriptomes for three major ocular tissues: cornea, retina, and RPE, using RNA-seq data from both adult and fetal tissues to create a high-quality pan-eye transcriptome. In addition to ocular samples, we used a subset of the GTEx data set to construct transcriptomes for tissues in 44 other locations across the body.

We used our gold-standard informed pan-eye *de novo* transcriptome to reveal hundreds of novel gene isoforms in the eye and analyze their potential impact on ocular biology and disease. We provide transcript annotation derived from our *de novo* transcriptomes as a resource to other researchers through an R package.

## Results

### Long-read PacBio RNA sequencing guides short-read *de novo* transcriptome construction

To evaluate the accuracy of short-read transcriptome construction, we first generated PacBio long-read RNA-seq data and Illumina short-read RNA-seq data from iPSC-RPE (Fig 1). These cells were differentiated using an optimized protocol, and thus minimal biological variation is expected (Blenkinsop et al., 2015), (Maruotti et al., 2015). We used these sequencing data to construct a long-read transcriptome and a short-read transcriptome. In our long-read transcriptome we found 1163239 distinct transcripts, and in our short-read transcriptome 366888 distinct transcripts

**Figure 1.**
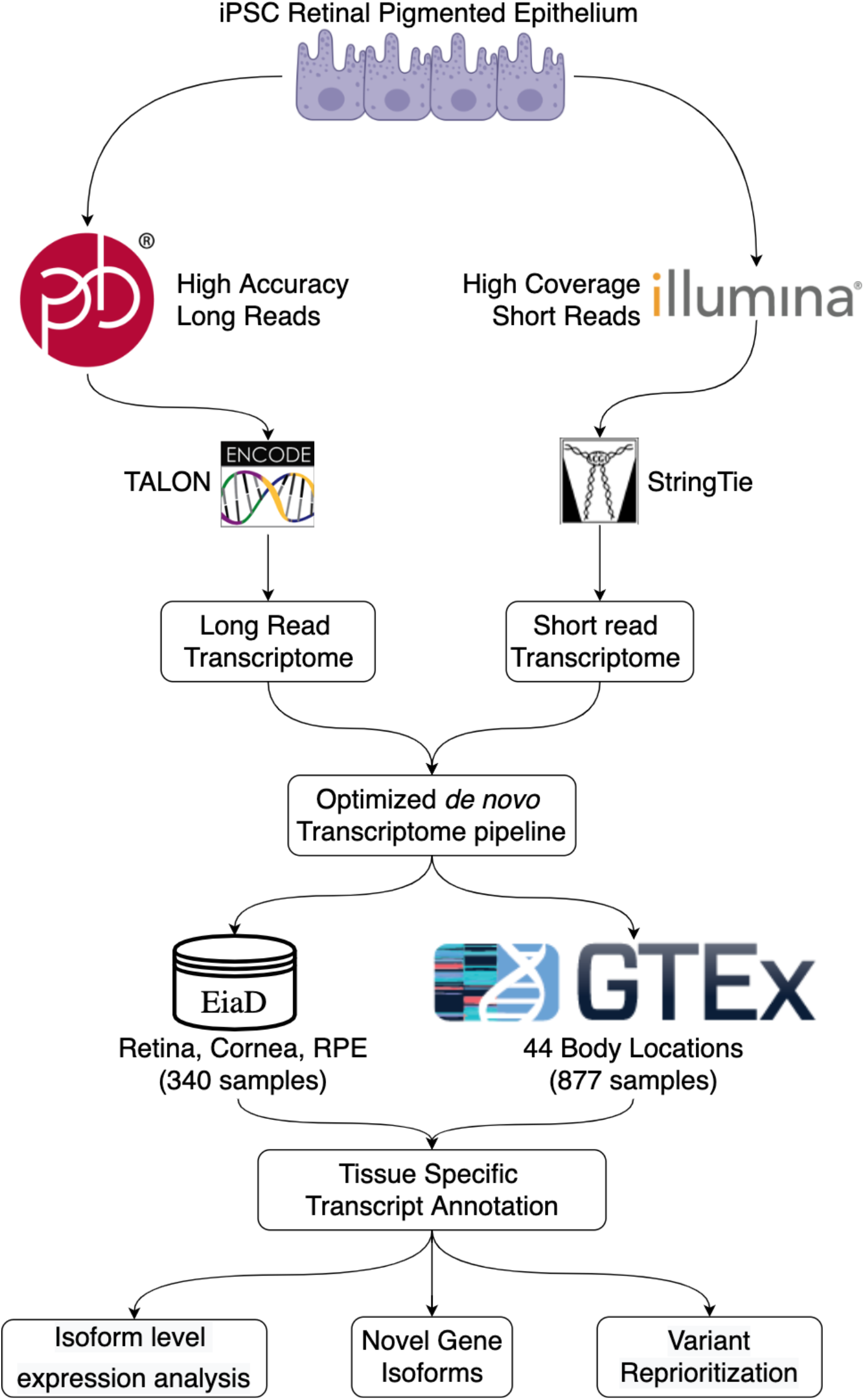
Workflow for long-read informed *de novo* transcriptome construction and analysis

In our initial comparison between short and long-read transcriptomes, we noticed a low transcriptome construction accuracy (see Methods) of 0.208. When we examined the transcript lengths of each build we saw that the two methods show very different transcript length distributions for both novel and previously annotated transcripts, with the short-read build was comprised mostly of smaller transcripts (Fig 2A). As the PacBio data was generated using two different libraries for 2000 bp and >3000 bp transcripts, we expected an enrichment for longer transcripts in the PacBio data set (Supplemental Figure 2). To assess accuracy relative to transcript length, we grouped transcripts by length in 1000 bp intervals, and compared accuracy between each group. We found that accuracy significantly improves for transcripts longer than 2000 bp. The construction accuracy is 0.426 and 0.137 for transcripts above and below 2000 bp, respectively.

**Figure 2.**
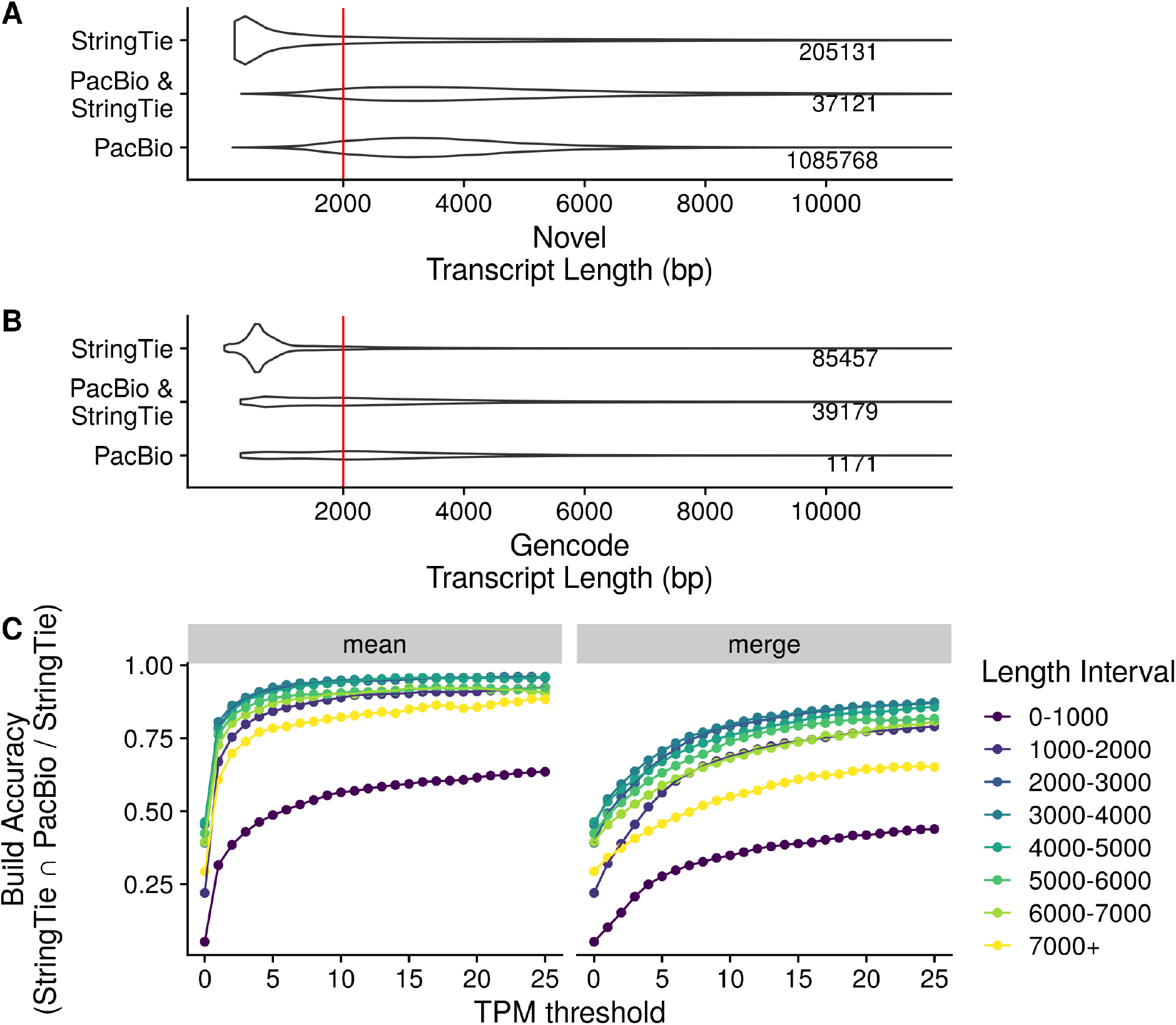
Transcript length and expression dictate transcriptome construction accuracy. A,B) Distributions of novel(A) and previously annotated(B) transcript lengths between PacBio (long-read) and Stringtie (short-read) transcriptomes. Each distribution is labeled with the total number of transcripts in the distribution C) short-read construction accuracy stratified by transcript length at different Transcripts Per Million (TPM)-based transcript exclusion thresholds. The “merge” method follows the protocol for constructing transcriptomes outlined by the StringTie authors and keeps any transcripts expressed above a specific TPM threshold in at least one samples. The “mean” method used by our pipeline keeps transcripts whose average expression across all samples is above a specific TPM threshold.

We experimented with various methods to remove spurious transcripts and improve construction accuracy. We first removed transcripts that were expressed <1 TPM in at least one sample as outlined in StringTie’s recommended protocol (Pertea et al., 2016). This improved construction accuracy to 0.475 for transcripts longer than 2000bp and 0.212 for transcripts shorter than 2000bp. As this accuracy was still fairly low, we tried different filtering schemes, including experimenting with machine learning-based strategies to identify transcripts that were computational artifacts (data not shown), but we found that the simplest approach with high performance was to retain transcripts that had an average TPM above a specific threshold(Fig 2C). In our downstream pipeline we keep transcripts that have at least an average of 1 TPM across all samples of the same subtissue type as this threshold achieved a build accuracy of 0.772 for transcripts longer than 2000Bp and retained 48470 transcripts within this short-read RPE dataset.

### Thousands of novel gene isoforms are detected in human subtissue-specific transcriptomes

We built transcriptomes from 340 publicly available ocular tissue RNA-seq samples curated in EiaD using an efficient Snakemake pipeline (Köster and Rahmann, 2012). We included both publicly collated non-disease, non-perturbed adult and fetal samples from cornea, retina, and RPE tissues, mined from 29 different studies (Table 1). Our fetal tissues consist of both human fetal tissues and human iPSC-derived tissue, as stem cell-derived tissue has been showed to closely resemble fetal tissue. We inlcude our iPSC-RPE samples originally used to develop our pipeline within this larger set of fetal RPE samples. (Klimanskaya et al., 2004). To more accurately determine the tissue specificity of novel ocular transcripts, we supplemented our ocular data set with 877 samples from 44 body locations across 22 major tissues from the GTEx project and constructed transcriptomes for each of these body locations (GTEx Consortium et al., 2017). We refer to each distinct body location as a subtissue here after.

**Table 1.**
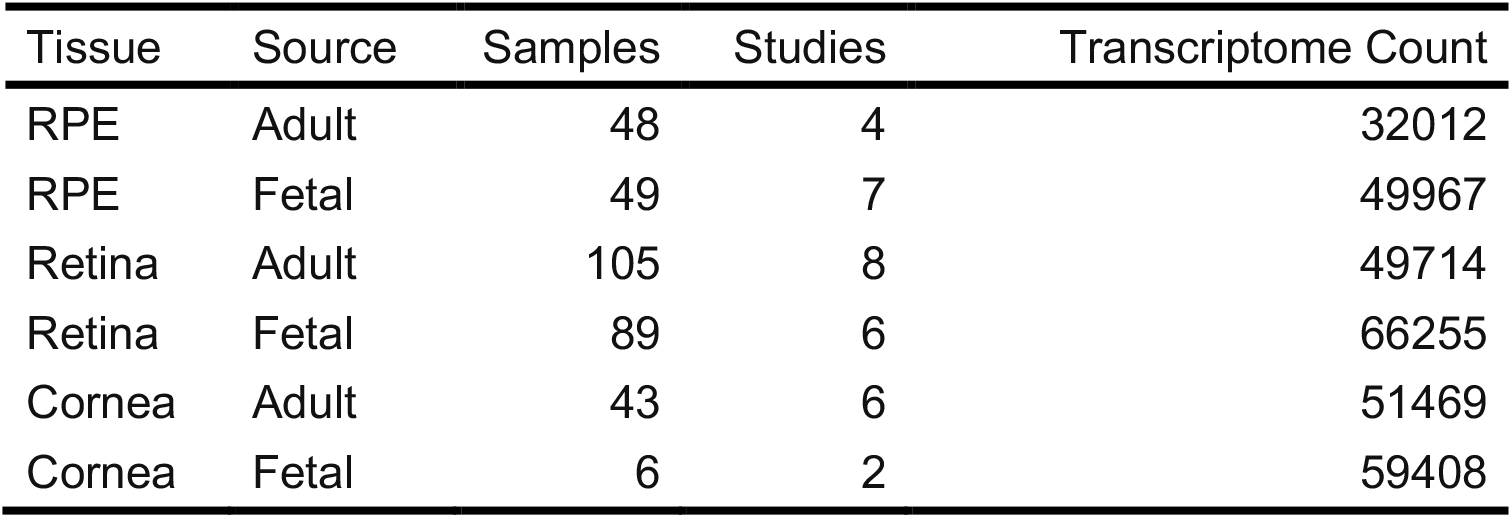
Ocular sample dataset overview and transcriptome count. Transcriptome count is defined as the number of unique transcripts expressed in a given tissue type

After initial construction of transcriptomes, we found 183442 previously annotated transcripts and 6241675 novel transcripts detected in at least one of our 1217 samples. We define a novel transcripts as all transcripts whose set of exons and introns do not exactly match that of an annotated transcript within the Gencode, Ensembl, UCSC, and Refseq annotation databases (Frankish et al., 2019), (Zerbino et al., 2018), (O’Leary et al., 2016). After using the filtering methods described above, we merged all subtissue specific transcriptomes into a single final transcriptome which contains 252983 distinct transcripts with 87592 previously annotated and 165391 novel transcripts, and includes 114.9 megabases of previously unannotated genomic sequence (Table 1). We refer to the final pan-body transcriptome as the DNTX annotation hereafter.

We split novel transcripts into two categories: novel isoforms, which are novel variations of known genes, and novel loci, which are previously unreported, entirely novel regions of transcribed sequence (Fig 3B). Novel isoforms are further classified by the novelty of their encoded protein: isoforms with novel open reading frame, novel isoforms with a known ORF, and isoforms with no ORF as noncoding isoforms (Fig 3A). The number of distinct ORFs was significantly less than the number of transcripts, with 43279 previously annotated ORFs and 46226 novel ORFs across all subtissues. Furthermore, across all subtissues there was an average of 10393 novel isoforms and 3716 novel ORFs.

**Figure 3.**
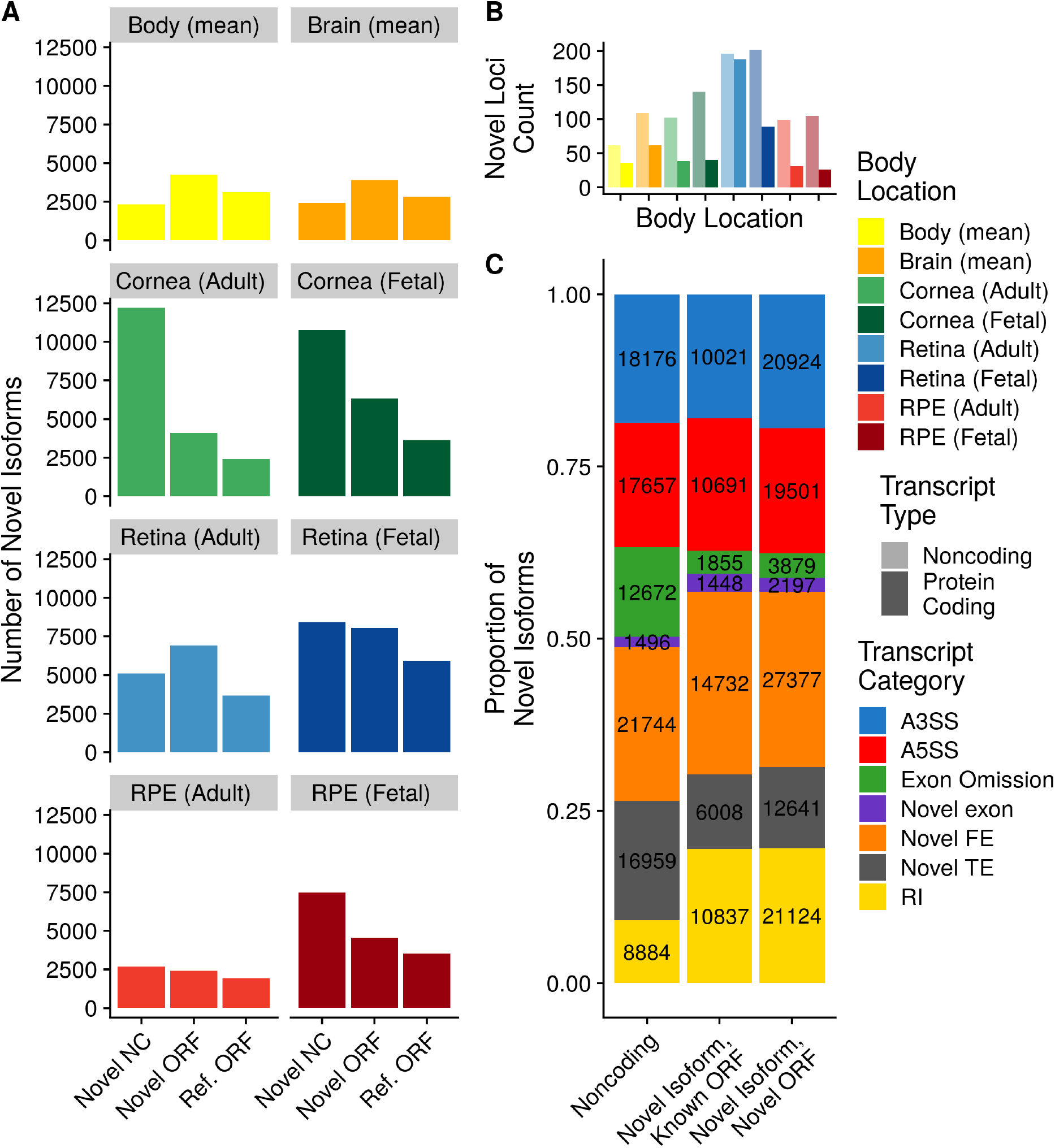
Overview of novel isoforms. A) Number of novel gene isoforms, grouped by transcript type. Brain and body represent an average of 13 and 34 distinct subtissues, respectively. B) Novel protein coding and noncoding loci. Novel exon composition of novel isoforms, by isoform type. Labels indicate number of transcripts. C) Classification of novel exon types, stratified by novel isoform type.

Novel isoforms can occur due to an omission of a previously annotated exon, commonly referred as exon skipping or the addition of an unannotated exon which we refer to as a novel exon. We further classified novel exons by the biological process that may be driving their formation: alternative promoter usage driving the addition of novel first exons (FE), alternative polyadenylation driving the addition of novel terminal exons (TE), and alternative splicing driving the formation of all novel exons that are not the first or last exon (Landry et al., 2003), (Tian and Manley, 2017), (Wang et al., 2015). We then split alternatively spliced exons into their commonly seen patterns, alternative 5’ splice site (A5SS), alternative 3’ splice site (A3SS), and retained introns (RI). Exons whose entire sequence was unannotated and is not a retained intron are fully novel exons. We note that all three of these mechanisms can lead to exon skipping, so for simplicity we grouped all novel isoforms resulting from exon skipping together. We found that the majority of novel exons within our dataset are novel FEs. We noticed that the majority of RI exons lead to novel ORFs, whereas novel isoforms with omitted exons more often lead to noncoding isoforms. (Fig 3C)

### *De novo* transcriptomes match previously published experimental data better than existing annotation

We validated *de novo* transcriptomes using three independent approaches. We first looked for evolutionary conservation since it is commonly accepted as a proxy for functional significance. We used the PhyloP 20 way species alignment, a measure of conservation between species, to calculate the average conservation score for each exon in the DNTX annotation and compared that to the average conservations score for each exon in the Gencode annotation (Pollard et al., 2010). We found that, on average, exons in the DNTX annotation are more conserved than exons in the Gencode annotation (pvalue <2.2e-16) (Supplemental Figure 2A).

Next, since we observed an enrichment in novel first and last exons within our data set, we decided to compare the TSS and TES within the DNTX annotation to two well-established annotation databases from FANTOM and the polyA Atlas (Noguchi et al., 2017), (Herrmann et al., 2020). We compared DNTX and Gencode TSS’s to CAGE-seq data from the FANTOM consortium; as CAGE-seq is optimized to detect the 5’ end of transcripts, we reasoned that it can serve as a valid ground truth set to evaluate TSS detection (Takahashi et al., 2012). We calculated the absolute distance of DNTX TSS’s to CAGE peaks, and compared them to the absolute distance of Gencode TSS’s to CAGE peaks. We found that, on average, DNTX TSS’s were closer to CAGE peaks than Gencode TSS’s (pvalue <2.2e-16)(Supplemental Figure 2B).

Finally, we evaluated TES’s using the polyA Atlas, which is comprised of polyadenylation signal annotation generated from aggregating 3’ seq data from multiple studies. As 3’-seq data is designed to accurately capture the 3’ ends of transcripts, it can similarly serve as a ground truth set to evaluate the accuracy of TES’s (Beck et al., 2010). We calculated the absolute distance of DNTX TES’s to annotated polyA signals and compared them to the absolute distance of Gencode TES’s to polyA signals. We found that on average DNTX TES’s are closer to annotated polyadenylation signals than gencode TSS’s (pvalue <2.2e-16) (Supplemental Figure 2C)

### *De novo* transcriptomes reduce overall transcriptome sizes

*De novo* transcriptomes removed on average 76.141 % of a subtissue’s base transcriptome. We defined base transcriptome for a subtissue as any transcript in the Gencode annotation with non-zero TPM in at least one sample of a given subtissue. This was a large reduction in transcriptome size and we wanted to ensure that we were not unduly discarding data. We quantified transcript expression of each sample using Salmon with two methods: once using the full gencode v28 human transcript annotation, and once using its associated subtissue specific transcriptome. We found that despite the 76.141 % reduction in number of transcripts between the base gencode and *de novo* transcriptomes (Supplemental Figure 3A), the per-sample Salmon mapping rate increased on average by 2.041 % indicating that the vast majority of gene expression data is retained within our transcriptome (Supplemental Figure 3B).

### Novel Isoforms are identified in ocular tissues

Using the pan-eye transcriptome, we compared the overlap in constructed novel isoforms across ocular subtissues and found that 77.968 % of novel isoforms are specific to a singular ocular subtissue (Fig 4A). Additionally, fetal-like tissues had more novel isoforms that their adult counterpart. For each novel isoform we then calculated fraction isoform usage (FIU), or the fraction of total gene expression a transcript contributes to its parent gene. We found that, on average, novel isoforms contributed to 20.584 % of their parent gene’s expression but in each subtissue we found multiple novel isoforms that contribute to the majority of their parent genes expression (Fig 4B)

**Figure 4.**
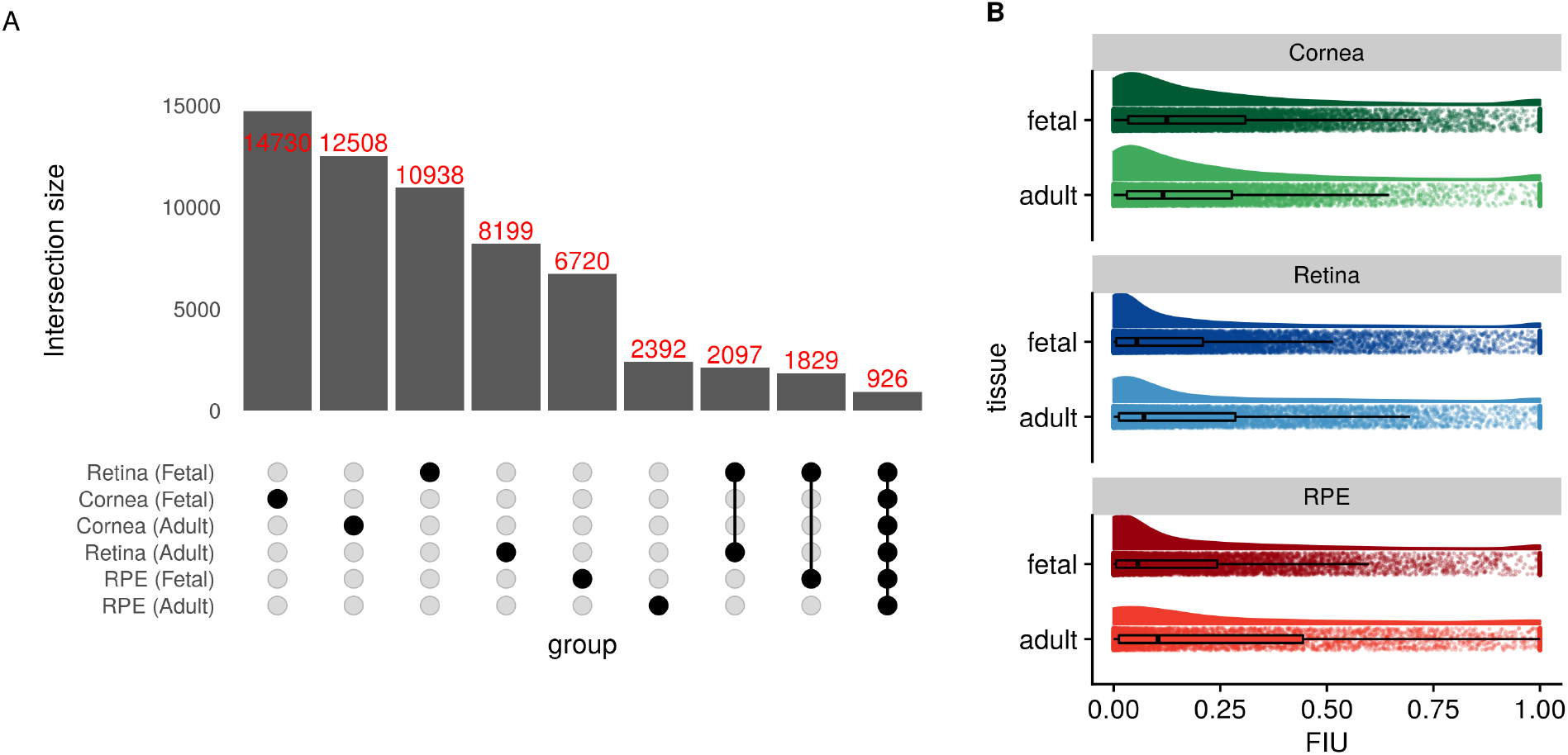
Overview of novel gene isoforms in the eye. A) Set intersection of novel isoforms in ocular transcriptomes. B) Boxplots of fraction isoform usage (FIU) overlaid over FIU data points with estimated distribution of data set above each boxplot.

### Differential usage of gene isoforms occurs during retinal development

Multiple studies have shown that gene isoforms play a significant role in eye development (Bharti et al., 2008), (Mellough et al., 2019). We hypothesized that the DNTX annotation provides additional insight into alternative isoform usage and identifies novel gene isoforms potentially involved in eye development. We used RNA-seq data of the developing retina from Mellough et al, an independent data set that we did not include for transcriptome construction, and used a subset of the DNTX annotation corresponding to fetal retina to quantify transcript expression and identify transcripts with significant changes in expression across retinal development. Transcripts that are differentially expressed (qvalue <.01) and have a mean FIU difference of .25 in at least one comparison of time points are indicative of differential transcript usage (DTU).

We analyzed 24 samples across 14 developmental days post fertilization and found 1717 transcripts across 812 genes displaying DTU (Fig 5A). We found that genes involved in DTU are enriched(qvalue <.05) for genes related to eye and neurological development (Fig 5B), and that hierarchical clustering of DTU transcripts generates an early stage and late stage cluster (Fig 5C). One of these genes, *MYO9A*, is a classical example of DTU. *MYO9A* is associated with the visual perception GO term, plays a role in ocular development, and has been associated with ocular disease (Gorman et al., 1999). While expression of *MYO9A* remains relatively unchanged across development, expression of two of its associated isoforms in fetal retina (Fig 5D) changes dramatically during development: a novel isoform is highly expressed early during development, but switched to the canonical isoform later in development (Fig 5E,F). This novel isoform contains a novel exon within the protein coding region of the isoform as well as novel last exon extending the 3’ UTR (Fig 5d). A full list of genes and transcripts displaying DTU is available in Supplemental Data (Supplemental Data 4).

**Figure 5.**
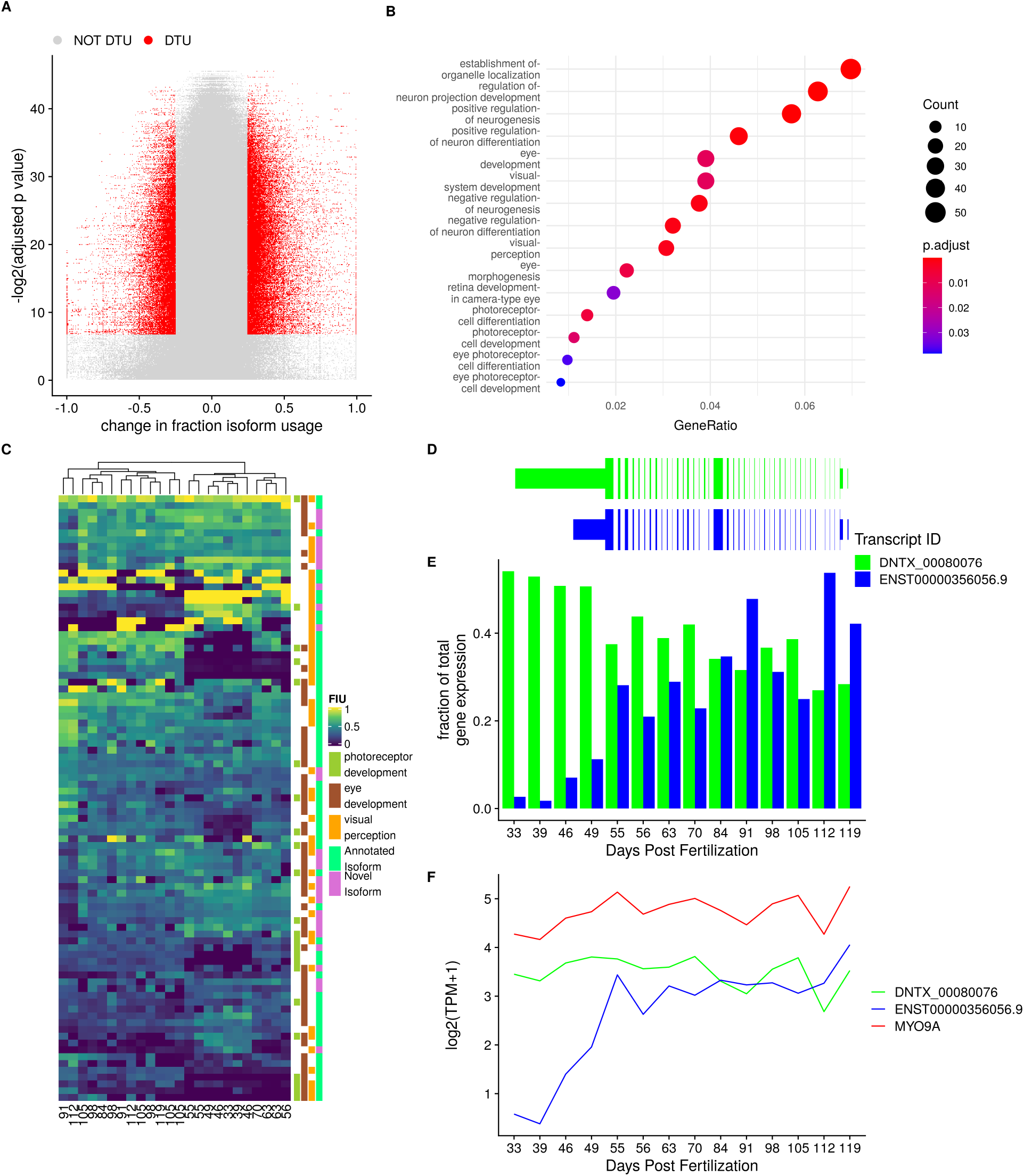
Differential Transcript usage during retinal development. A) Volcano plot of tested transcripts B) Dot plot for gene set enrichment analysis C) Heatmap of hiearchical clustering of transcripts with DTU associated with eye development D) Transcript models for *MYO9A*, a gene undergoing DTU E) FIU change in *MYO9A* FIU across development F) average log-transformed TPM expression of *MYO9A* across retinal development

### *De novo* transcriptomes allow for a more precise variant prioritization

The identification of a disease-causing variant through genome sequencing is a common step in diagnosing genetic disease, when disease causing variants cannot be determined from exonic sequencing. Prediction of a variant’s biological impact and subsequent variant prioritization is a fundamental step in this process. Many methods for predicting variant effects on protein function or gene expression are based on location within the body of a transcript; for example variants that disrupt splice sites and start/stop codons are considered to be the most damaging, while variants within intronic and intergenic regions have unknown impact or are not classified, and, thus, are not included for further consideration. However, multiple studies have identified pathogenic deep intronic variants for retinal dystrophies (Braun et al., 2013), (Bauwens et al., 2019), (Zernant et al., 2014), (Sangermano et al., 2019), (Jamshidi et al., 2019), (Mayer et al., 2016), (Geoffroy et al., 2018). Pathogenic intronic variants are thought to function by introducing a novel splice site, disrupting regulatory motifs, or altering a tissue-specific transcript. To explore this third possibility, we mapped known pathogenic intronic variants onto novel isoforms within the *de novo* transcriptomes.

We used a list of 129 intronic and noncoding variants previously identified as pathogenic for a retinal dystrophy and predicted the effect of these variants with Ensembl’s Variant Effect Predictor using a subset of the DNTX annotation corresponding to fetal and adult retina as the input transcript annotation. We identified ten variants whose predicted effect increased in severity due the presence of a novel gene isoform in a previously intronic region (Table 2). Seven of these variants were in deep intronic hotpsots known for pathogenic variation within the gene ABCA4.

**Table 2.** Pathogenic variants previously considered intronic that are on expressed transcripts in the retina *de novo* transcriptome. Canonical human genome variation society (HGVS) annotation is based on transcripts from the RefSeq annnotation. Predicted consequences were generaed with the Variant Effect Predictor(VEP)

| Gene Name | Associated Disease | Location (hg19) | Canonical Variant HGVS | Gencode Predicted Consequence | DNTX Predicted Consequence | Published Study |
| --- | --- | --- | --- | --- | --- | --- |
| ABCA4 | ABCA4-associated maculopathy | Chr1:94481967 C>T | c.5197-557G>T, NM_000350.2 | intron variant, downstream gene variant | 5 prime UTR variant | Bauwens et al. |
|  |  | Chr1:94546814 G>C | c.859-540C>G, NM_000350.2 | intron variant | non coding transcript exon variant |  |
|  | Stargardt disease | Chr1:94484001 C>T | c.5196+1137G>A, NM_000350.2 | intron variant, downstream gene variant | 5 prime UTR variant | Braun et al.<br>Zernant et al. |
|  |  | Chr1:94484082 T>G | c.5196+1056A>G, NM_000350.2 | intron variant, downstream gene variant | 5 prime UTR variant |  |
|  |  | Chr1:94526934 T>G | c.1938-619A>G, NM_000350.2 | intron variant, splice region variant, non coding transcript variant | non coding transcript exon variant | Zernant et al. |
|  |  | Chr1:94527698 G>C | c.1937+435C>G, NM_000350.2 | intron variant, upstream gene variant | non coding transcript exon variant | Sangermano et al. |
|  |  | Chr1:94546780 C>G | c.859-506G>C, NM_000350.2 | intron variant | non coding transcript exon variant |  |
| IFT140 | Ciliopathy | Chr16:1576595 C>A | c.2577+25G>A, NM_014714.3 | upstream gene variant, intron variant, NMD transcript variant, non coding transcript exon variant, non coding transcript variant | missense variant | Geoffroy et al. |
| PROM1 | Cone-rod dystrophy | Chr4:15989860 T>G | c.2077-521A>G, NM_006017.2 | intron variant, upstream gene variant | 5 prime UTR variant | Mayer et al. |
| RPGRIP1 | RPGRIP1-mediated inherited retinal degeneration | Chr14:21789588 G>A | c.1611+27G>A, NM_020366.3 | intron variant, non coding transcript variant, upstream gene variant, synonymous variant, NMD transcript variant, downstream gene variant | 5 prime UTR variant | Jamshidi et al. |

These variants were spanned by three distinct novel isoforms with two containing open reading frames (ORFs) encoding only the carboxy-terminus of the canonical protein isoform, and one noncoding spanning the proximal half of the canonical isoform (Fig 6). *ABCA4* expression and function has also been observed in RPE (Lenis et al., 2018). However, we did not observe these transcripts in RPE, suggesting that these pathogenic variants are primarily affecting retinal-specific *ABCA4* transcripts. We note that these transcripts have not been experimentally validated.

**Figure 6.**
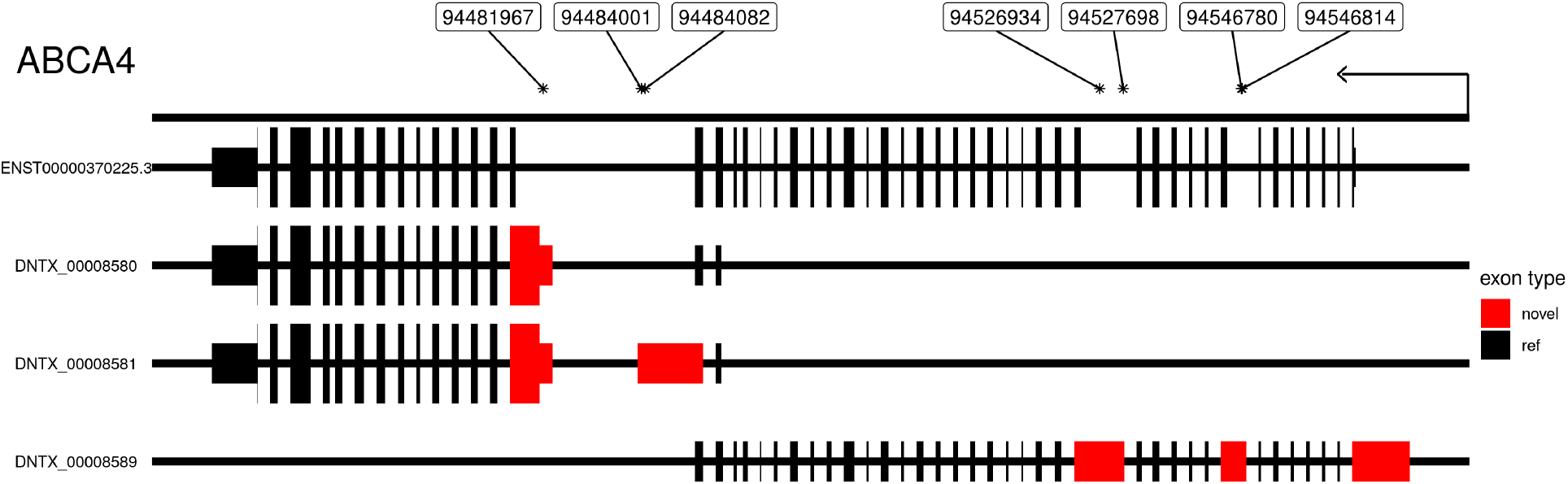
Transcript models for selected Isoforms of *ABCA4* along with location of pathogenic intronic variants. Location is on the hg19 human genome build. Thick lines indicate protein coding regions. Arrow indicates direction of transcription. Introns not drawn to scale

To further highlight the potential importance of *de novo* transcriptomes for future genetic tests we determined how many genes associated with retinal disease from RetNet have novel isoforms (sph.uth.edu/retnet/). We found that within the set of genes with novel isoforms, there is significant enrichment of retinal disease genes (hypergeometric pvalue = 3.4e-04), with 220 out of 379 RetNet genes having a novel isoform. A full list of these genes is available in the Supplementary data(supplemental data 5).

### A companion visualization tool enables easy use of *de novo* transcriptomes

To make our results easily accessible we designed a R-Shiny app for visualizing and accessing our *de novo* transcriptomes. For each subtissue we show the FIU for each transcript associated with a gene (Fig 7A). We show the exon-intron structure of each transcript and mousing over exons show genomic location overlapping SNPs, and phylogenetic conservation score (Fig 7B). We additionally show a barplot of the fraction of samples each transcript was constructed in (Fig 7C). Users can also download the *de novo* transcriptomes for selected subtissues in GTF and fasta format. Instructions to download and run the app are available at https://github.com/vinay-swamy/ocular_transcriptomes_shiny. While visualization of direct transcript expresion is not a part of this app, it can be viewed in the eyeIntegration app (Swamy and McGaughey, 2019) by selected ‘DNTX’ as the transcript annotation. Finally, we provide all code as a Snakemake workflow and provide a Docker container with all software required for the pipeline available at https://github.com/vinay-swamy/ocular_transcriptomes_pipeline

**Figure 7.**
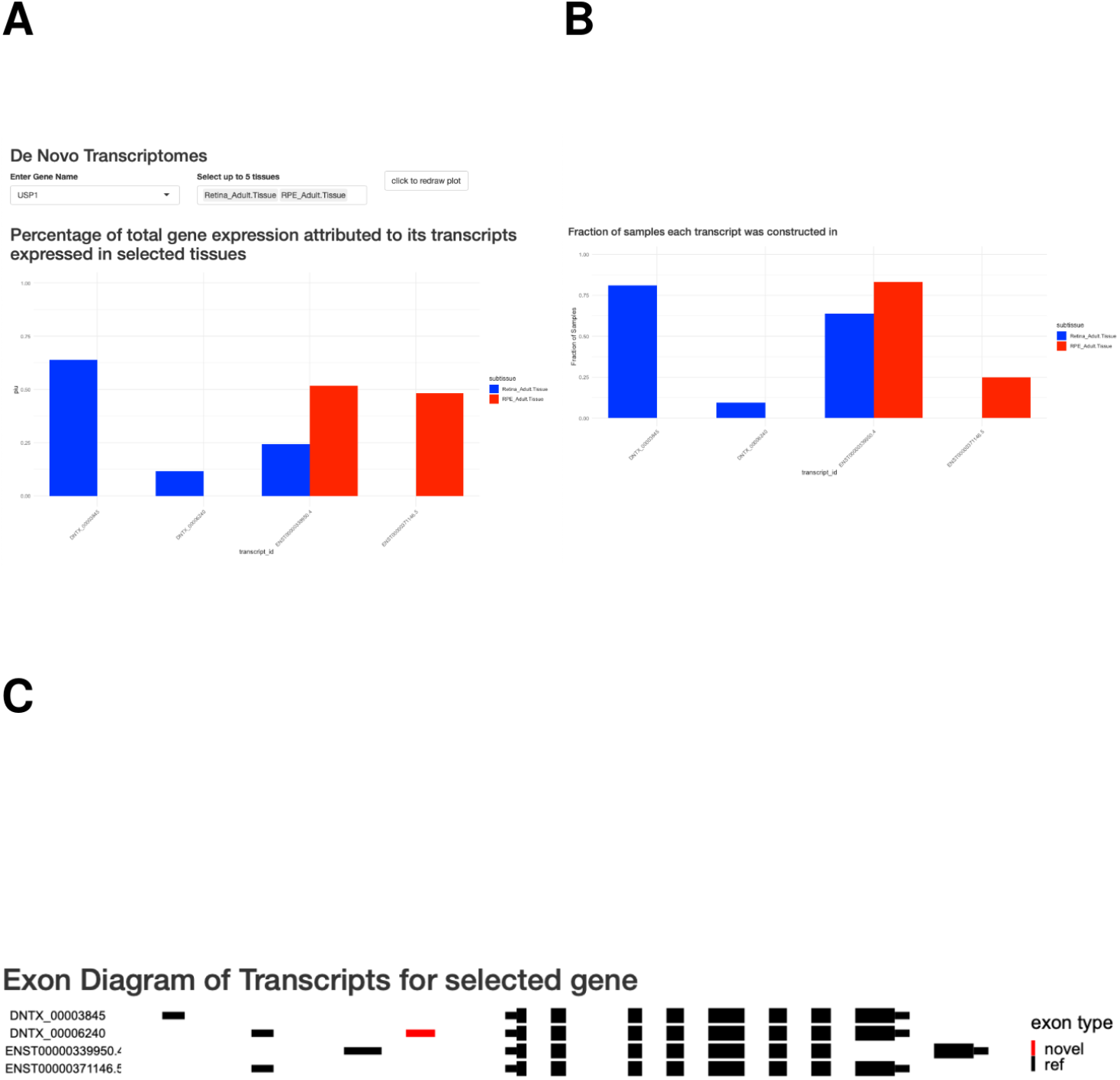
Screenshots from dynamic *de novo* transcriptome visualization tool. A). FIU bar plot for selected gene and subtissue. B). Exon level diagram of transcript body Thicklines represent coding region of transcript. novel exons colored in red. Tooltip contains genomic location and phylop score C) Bargraph of fraction of samples within dataset each transcript was consructed in by tissue.

## Discussion

Motivated by the lack of a comprehensive transcriptome for the eye, we constructed transcriptomes for adult and fetal retina, RPE and cornea. By using long-read RNA-seq data to calibrate our short-read construction pipeline, we were able to identify biologically relevant transcriptomes. We found that concordance between long and short-read-based transcriptomes is directly related to transcript length and transcript expression. We saw a clear inability within the PacBio data set to accurately detect transcripts shorter than 2000bp for both previously annotated and novel transcripts. As many of the transcripts constructed using short-reads are below this threshold, long-read sequencing data enriched for smaller transcript sizes would provide greater insight in future studies.

We used a large dataset compiled from published RNA-seq data to build the pan-eye transcriptomes, an approach that has several key advantages. First, the large sample size overcomes the noisy nature of RNA-seq data. Second, as the cohort is constructed from many independent studies, we are more confident that the transcriptomes accurately reflect the biology of their originating subtissue and are not a technical artifact due to preparation of the samples. As another line of evidence, the *de novo* transcriptomes match existing large scale data sets and are more conserved than existing annotations (Supplemental Figure 2).

In each ocular subtissue we examined, we found hundreds of novel gene isoforms, many of which were novel due to novel exons. Within ocular subtissues, these novel isoforms are most commonly specific to single subtissue. This makes sense as a majority of the exons in our *de novo* transcriptomes are first and last exons, which have been previously shown to significantly contribute to the tissue specificity of gene isoforms (Reyes and Huber, 2018). We also found that on average novel isoforms represent about 20.584 % of their parent gene’s expression. Future studies are needed to identify the function of these isoforms. One possibility is that some of these isoforms are only expressed in rare cell types, as transcript annotation was previously shown to be incomplete in rare cell types (Zhang et al., 2020). This especially makes sense in the retina which contain over a dozen distinct cell types, several of which contribute to 5% or less of the total cell population (Yan et al., 2020). As we imposed a strict expression filter as part of our transcriptome pipeline, we may have removed transcripts specific to rare cell types.

In conclusion, we created the first pan-eye transcriptome annotation and showed that it is useful in understanding the role of gene isoforms in ocular biology and improving the ability to diagnose inherited eye diseases. We hope this work is useful as a starting point for other researchers; [delete] to make the transcriptomes easily accessible to other researchers we designed a webapp both for visualization and to quickly access tissue-specific annotation files. We believe this project will enable other researchers to explore new research directions and answer long pending questions.

## Methods

### Generation of PacBio long-read RNA sequencing data and Illumina short-read RNA sequencing data

Human iPSCs were differentiated into RPE using previously described protocols in (Bryan et al., 2018) and (May-Simera et al., 2018). iPSC-derived RPE (iPSC-RPE) cells at 42 days post differentiation were lysed with TRIzol reagent (Thermo Fisher Scientific; cat # 15596026) and total RNA was isolated using the Direct-zol RNA MiniPrep Kit (Zymo Research, Irvine, CA). 5-6 µg total RNA that passed quality control metric (RIN >.9) were used for PacBio library preparation. For PacBio HiFi circular consensus sequencing(CCS), libraries were prepared following the “Procedure-Checklist-Iso-Seq-Express-Template-Preparation-for-Sequel-and-Sequel-II-Systems” protocol. Two libraries were generated: one to capture transcripts 2 kilobases(kb) or smaller, and one to capture transcripts between 2-5kb. Sequencing was done on the PacBio Sequel II system for a movie time of 24 hours.

For Illumina sequencing, Poly-A selected stranded mRNA libraries were constructed from 0.5-1 µg total RNA using the Illumina TruSeq Stranded mRNA Sample Prep Kits according to manufacturer’s instructions. Amplification was performed using 10-12 cycles to minimize the risk of over-amplification. Unique dual-indexed barcode adapters were applied to each library. Libraries were pooled in equimolar ratio and sequenced together on a HiSeq 4000. At least 57 million 75-base read pairs were generated for each individual library. Data was processed using illumina Real Time Analysis (RTA) version 2.7.7. All library preparation and sequencing was performed at the National Institutes of Health Intramural Sequencing Center (NISC).

### Code availability and software versions

To improve reproducibility, all code used for both the analyzing the data and generating the figures for this paper was written as multiple Snakemake pipelines. Each Snakefile contains the exact parameters for all tools and scripts used in each analysis. (Köster and Rahmann, 2012) All code (and versions) used for this project is publicly available in the following github repositories: https://github.com/vinay-swamy/ocular_transcriptomes_pipeline (main pipeline), https://github.com/vinay-swamy/ocular_transcriptomes_longread_analysis (long-read analysis pipeline), https://github.com/vinay-swamy/ocular_transcriptomes_paper (figures and tables for this paper), https://github.com/vinay-swamy/ocular_transcriptomes_shiny (webapp). Additionally, all Snakefiles are included as supplementary data.(supplementary data files 1-3)

### Analysis of long-read data

PacBio sequencing movies were processed into full length, non-chimeric (FLNC) reads using the IsoSeq3 3.1.2 pipeline in the PacBio SMRT link v7.0 software. The existing ENCODE long-read RNA-seq pipeline (https://github.com/ENCODE-DCC/long-read-rna-pipeline) was rewritten as a Snakemake workflow as follows. Transcripts were aligned to the human genome using minimap2(18), using an alignment index built on the gencode v28 primary human genome. Sequencing errors in aligned long-reads were corrected using TranscriptClean (19) with default parameters. Splice junctions for TranscriptClean were obtained using the TranscriptClean accessory script “get_SJs_from_gtf.py” using the gencode v28 comprehensive transcript annotation as the input. A list of common variants to avoid correcting were obtained from the ENCODE portal (https://www.encodeproject.org/files/ENCFF911UGW/). The long-read transcriptome annotation was generated with TALON (20). A TALON database was generated using the talon_initialize_database command, with all default parameters, except for the “–5P” and “– 3p” parameters. These parameters represent the maximum distance between close 5’ start and 3’ ends of similar transcript to merge and were both set to 100 to match parameters used in later tools. Annotation in GTF format was generated using the talon_create_GTF command, and transcript abundance values were generated using the talon_abundance command.

### Analysis of short-read RPE data

Each sample was aligned to the Gencode release 28 hg38 human genome assembly using the genomic aligner STAR and the resulting BAM files were sorted using samtools sort (Frankish et al., 2019),(Dobin et al., 2013),(Li et al., 2009). For each sorted BAM file, a per-sample base transcriptome was constructed using StringTie with the Gencode v28 comprehensive annotation as a guiding annotation (Frankish et al., 2019),(Pertea et al., 2015). All sample transcriptomes were merged with the long-read transcriptome using gffcompare(Pertea and Pertea, 2020) with default parameters. We note that the default values for the distance to merge similar 5’ starts and 3 ends of transcripts in gffcompare is the same to what we chose for TALON. We defined the metric construction accuracy, used to evaluate short-read transcriptome construction as the following:

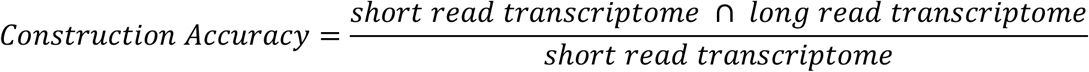

### Construction of subtissue-specific transcriptomes

We constructed transcriptomes for 1217 samples in the Eye in a Disk(EiaD), a dataset generated from aggregating publically available healthy, unperturbed RNA-seq samples from 50 distinct locations of the body across 29 different studies. Specific information on how this dataset was generated is detailed in the methods from our previous work (Swamy and McGaughey, 2019). We constructed a transcriptome for each sample, and merged samples together to create 50 subtissue-specific transcriptomes. We define subtissue as a unique body location and are either temporally different versions of the same tissue(adult vs fetal tissue), or different regions of a larger tissue (cortex vs cerebellum in brain). Tissue refers to complete whole tissue (retina, brain, liver). For each subtissue-specific transcriptome, we removed transcripts that had an average expression less than 1 Transcripts Per Million (TPM) across all samples of the same subtissue type. All subtissue-specific transcriptomes were merged to form a single unified annotation file in general transfer format(GTF) to ensure transcript identifiers were the same across subtissues. We merged all ocular subtissue transcriptomes to generate a separate pan-eye transcriptome.

### Subtissue specific transcriptome quantification

For each resulting subtissue specific transcriptome, we extracted transcript sequences using the tool gffread and used these sequences to build a subtissue-specific quantification index using the index mode of the alignment-free quantification tool Salmon (Pertea and Pertea, 2020), (Patro et al., 2017). For each sample, we quantified transcript expression using the quant mode of Salmon, using a sample’s respective subtissue specific quantification index. We similarly quantified all ocular samples using the pan-eye transcriptome and the Gencode v28 reference transcriptome.

### Annotation of novel exons

First, a comprehensive set of distinct, annotated exons was generated by merging exon annotation from gencode, ensembl, UCSC, and refseq. We then defined a novel exon as any exon within our transcriptomes that does not exactly match the chromosome, start, end and strand of an annotated exon. Novels exons were classified by splitting exons into 3 categories: first, last, and middle exons. We then extracted all annotated exon start and stop sites from our set of previously annotated exons. Novel middle exons that have an annotated start but an unannotated end were categorized as a novel alternative 3’ end exons and similarly novel middle exons with an unannotated start but annotated end were categorized as a novel alternative 5’ start exons. Novel middle exons whose start and end match annotated exon start and ends were considered retained introns. Novel middle exons whose start and end do not match annotated starts and ends were considered fully novel exons. We then classified novel first and last exons. Novel first exons were first exons whose start is not in the set of annotated exon starts, and novel last exons were terminal exons whose end is not in the set of annotated exon ends. This analysis of novel transcripts is implemented in our Rscript “annotate_and_make_tissue_gtfs.R”.

### Validation of DNTX with phylop, CAGE data, and polyA signals

PhyloP scores for the phylop 20-way multi species alignment were downloaded from UCSC’s FTP server on October 16th, 2019 and converted from bigWig format to bed format using the wig2bed tool in BEDOPs (Pollard et al., 2010), (Neph et al., 2012). The average score per exon in both the gencode and DNTX annotation was calculated by intersecting exon locations with phylop scores and then averaging the per base score for each exon, using the intersect and groupby tools from the bedtools suite, respectively. Significant difference in mean phylop score was tested with a Mann Whitney U test.

CAGE peaks were download from the FANTOM FTP server (https://fantom.gsc.riken.jp/5/datafiles/reprocessed/hg38_latest/extra/CAGE_peaks/hg38_fair+new_CAGE_peaks_phase1and2.bed.gz) on June 15th 2020 (Noguchi et al., 2017). Transcriptional start sites (TSS) were extracted from gencode and DNTX annotations; TSS is defined as the start of the first exon of a transcript. Distance to CAGE peaks was calculated using the closest tool in the bedtools suite. Significant difference in mean distance to CAGE peak between DNTX and gencode annotation was tested with a Mann Whitney U test.

Polyadenylation signal annotations were downloaded from the polyA site atlas (https://polyasite.unibas.ch/download/atlas/2.0/GRCh38.96/atlas.clusters.2.0.GRCh38.96.bed.gz) on June 15th 2020 (Herrmann et al., 2020). Transcriptional end sites(TES) were extracted from gencode and DNTX annotations; TES is defined as the end of the terminal exon of a transcript. Distance to polyA signal was calculated using the closest tool in the bedtools suite (Quinlan and Hall, 2010). Significant difference in mean distance to polyA signal was tested with a Mann Whitney U test.

### Identification of novel protein coding transcripts

Protein-coding transcripts in the unified transcriptome were identified using the TransDecoder suite (Haas et al., 2013). Transcript sequences in fasta format were extracted from the final pan-body transcriptome using the TransDecoder util script “gtf_genome_to_cdna_fasta.pl”. Potential open reading frames(ORFs) were generated from transcript sequences using the LongestORF module within TransDecoder, and the single best ORF for each transcript was extracted with the Predict module within Transdecoder. The resulting ORFs were mapped to genomic locations with the TransDecoder util script “gtf_to_alignment_gff3.pl”. For each ORF start and stop codons were extracted with the script “agat_sp_add_start_stop.pl” scripts from the AGAT toolkit (https://github.com/NBISweden/AGAT/). Transcripts with no detectable ORF or missing a start or stop codon were labelled as noncoding.

### Analysis of novel isoforms in eye tissues

An Upset plot was generated using the ComplexUpset package (https://github.com/krassowski/complex-upset) (Lex et al., 2014). Fraction Isoform Usage (FIU) was calculated for each transcript *t* associated with a parent gene *g* using the following formula: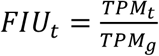. Raincloud plots of FIU were generated using the “R_Rainclouds” R package (Allen et al., 2019).

### Analysis of fetal retina RNA-seq data

RNA-seq samples from Mellough et al. were obtained from EiaD, and were not included in the main dataset used for building transcriptomes. Outliers within the dataset were identified by first performing principal component analysis of transcript level expression data, calculating the center of all data using the first two principal components, and subsequently removing five samples furthest away from the center of all data. The remaining samples were normalized using calcNormFactors from the R package edgeR and converted to weights using the voom function from the R package limma (Robinson et al., 2010), (Ritchie et al., 2015). Differential expression was modeled using the lmFit function using developmental time point as the model design and tested for significant change in expression using the Ebayes function from limma. Gene Set enrichment was tested using the R package clusterprofileR (Yu et al., 2012). Heatmaps were generated using the ComplexHeatmap package (Gu et al., 2016).

### Prediction of variant impact using *de novo* transcriptomes

Noncoding variants previously associated with retinal disease from the Blueprint Genetics Retinal dystrophy panel were obtained from the Blueprint Genetics website (https://blueprintgenetics.com/tests/panels/ophthalmology/retinal-dystrophy-panel/). The variants were converted from HGVS to VCF format using a custom python script “HGVS_to_VCF.py”. This VCF was then remapped to the hg38 human genome build using the tool crossmap (Zhao et al., 2014). The VCF of variants was used as the input variants for the Variant Effect Predictor(VEP) tool from Ensembl, with each subtissue specific transcriptome as the input annotation (McLaren et al., 2016). VEP was additionally run using the gencode v28 comprehensive annotation as the input annotation to identify variants whose predicted impact increased in severity.

### Figures, Tables, and Computing Resources

All statistical analyses, figures and tables in this paper were generated using the R programming language. (R Core Team, 2019) A full list of packages and versions can be found in the supplementary file session_info.txt. All computation was performed on the National Institutes of Health high performance computer system Biowulf (hpc.nih.gov).

## Competing Interests

All authors declare no Competing interests.

## Supplemental Figures

**Supplemental Figure 1.**
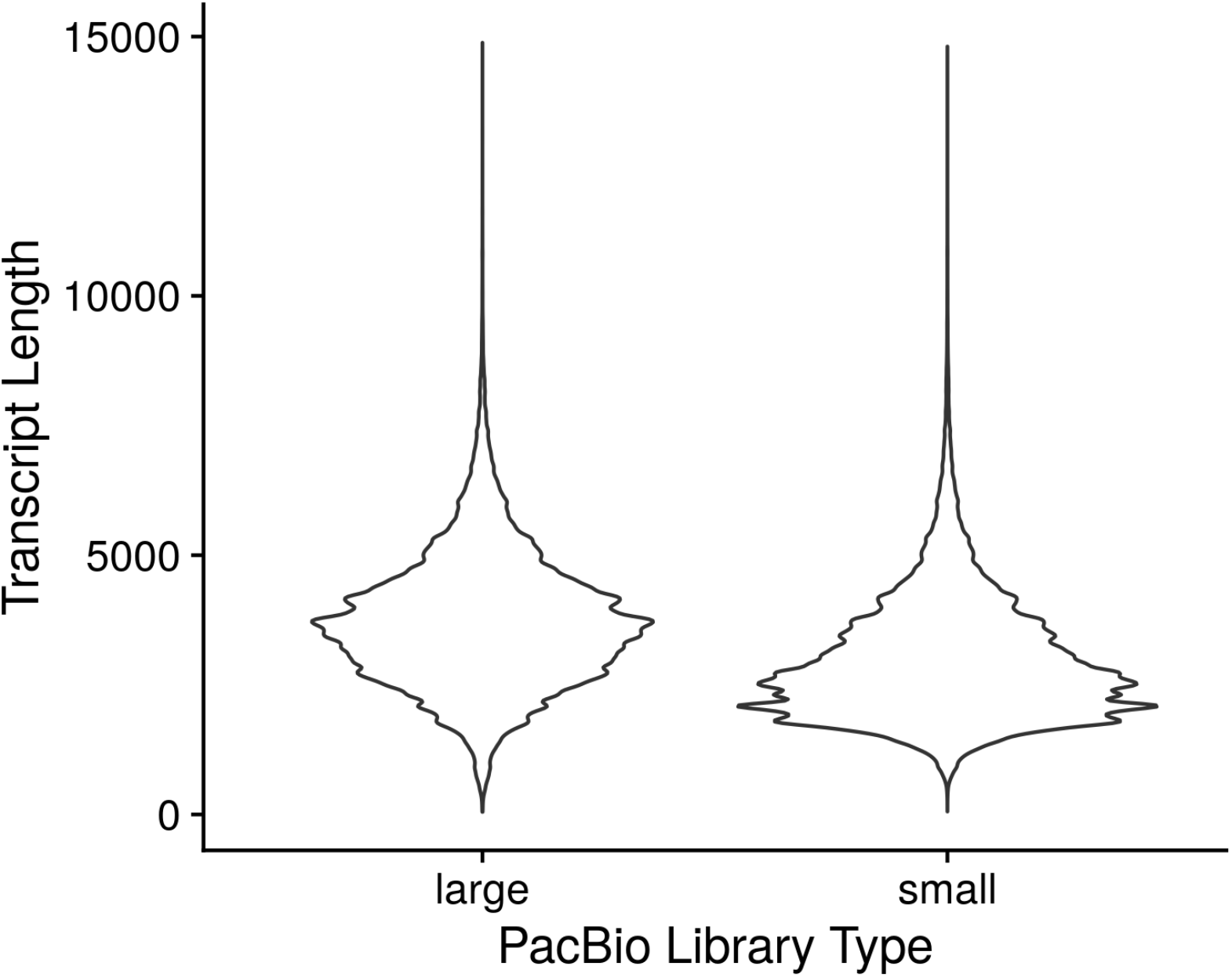
Distribution of PacBio long-read lengths for two library sizes.

**Supplemental Figure 2.**
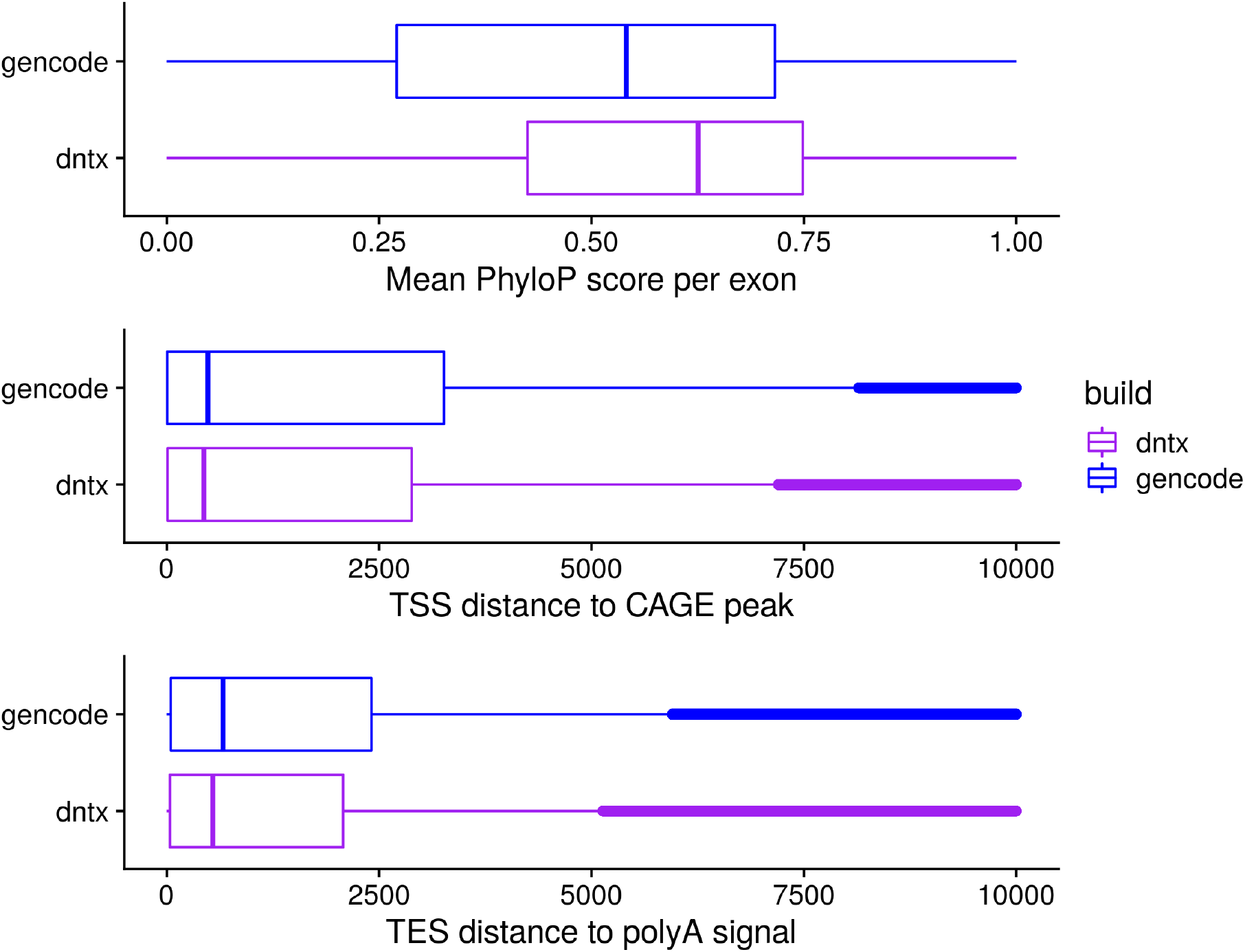
Comparison of DNTX annotation to Gencode annotation. A) Average per exon Phylop score for Gencode and DNTX transcripts. B) Average distance of DNTX transcriptional start sites (TSS) and Gencode TSS to CAGE-seq peaks from the FANTOM consortium. C) Average distance of DNTX transcriptional end sites (TES) and Gencode TES to polyadenylation signals in the PolyA site atlas.

**Supplemental Figure 3.**
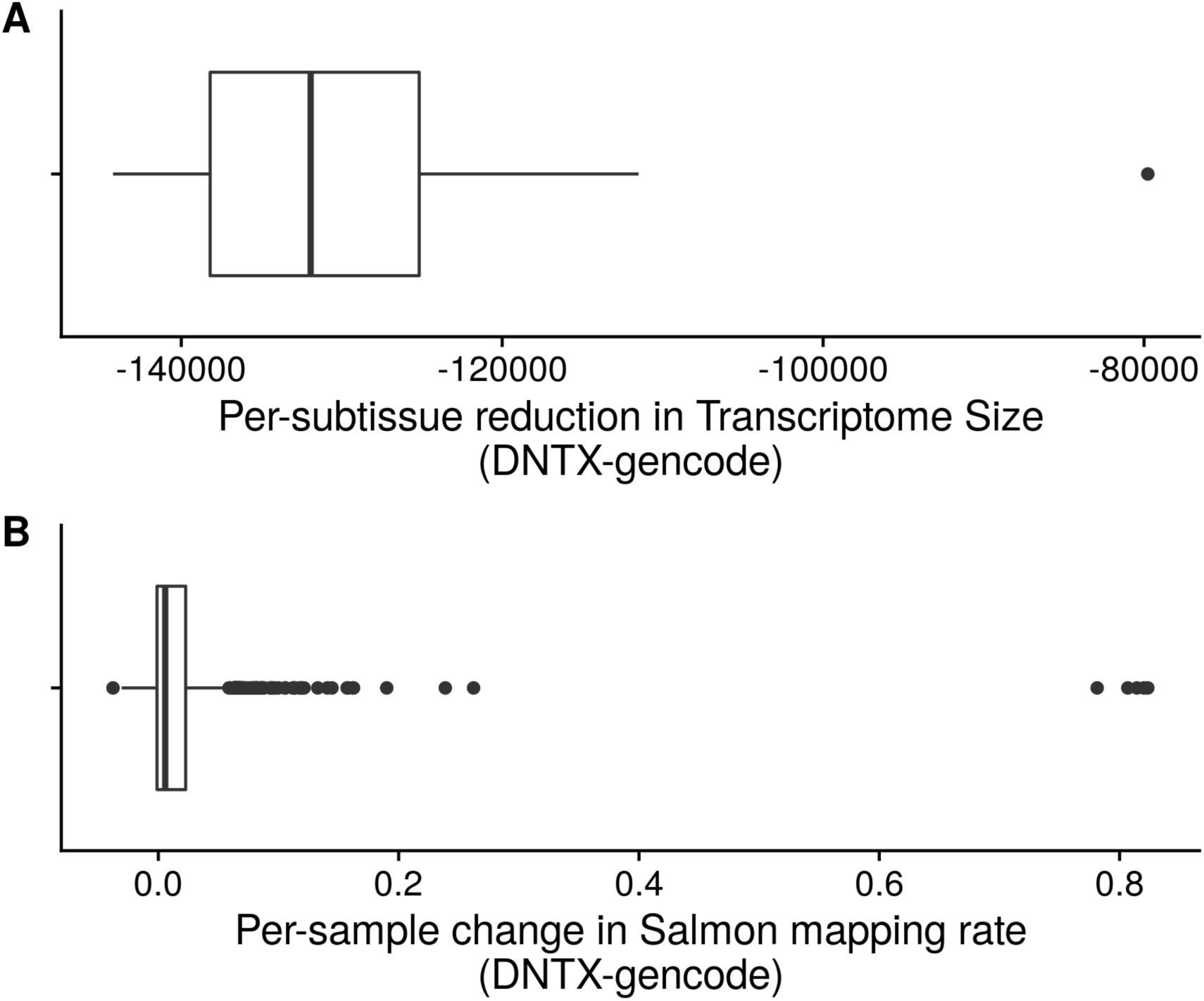
Comparison of Salmon mapping rate change vs transcriptome size decrease.

## Notes

https://github.com/vinay-swamy/ocular_transcriptomes_paper

https://github.com/vinay-swamy/ocular_transcriptomes_shiny

https://github.com/vinay-swamy/ocular_transcriptomes_pipeline

https://github.com/vinay-swamy/ocular_transcriptomes_longread_analysis

### Competing Interest Statement

The authors have declared no competing interest.

### Summary of Updates

The figure 2 Caption was was expanded for clarity; The citation format was changed; The abstract and introduction were slightly modified to remove areas of ambiguity. Changed title to reduce character count for journal submission

