## Supplemental Data 1 for "A long read optimized *de novo* transcriptome pipeline reveals novel ocular developmentally regulated gene isoforms and disease targets": 261644_file03

# pylint: skip-file
'''
config file:
    sampleFile: tab seperated file  containing sample info
    refFasta_url: link to refernece fastq_path
    salmon_version:
    sratoolkit_version:
notes:
-the 5.2 version requires specifying directorys in output section of rule iwth directory(). Biowulf currently using 5.1
-need to make a rule to download all Gencode refs

11/05/19
    REevesied 
-09/20/19
    Revised strategy: merge w/o genome guide with 1 TPM as c/o. quantify with salmon, remove tx with avg count< 1, and high
    bootstrap variance. filter removed transcripts from gtf, and then requantify. and then merge gtfs together.
06/10/19 changes
- I added way to much downstream stuff to the pipeline, assuming that the splicing/trnascriptome stuff was more accurate
  than it actually was, so I removed a lot stuff,  hmmer, converting to b37, getting a gff3, to focus more on improving
  the accuracy of the exon detections
- using the comprehensive gencode annotation for everything
- no longer realigning to stringtie gtf, this caused a lot of problems, an it makes a little more sense over all now,
  as both tools are blind to each other
- strategy for imporving - use a model trained on exons that get longer/shorter present in reference annotation
    - a model is trained for each sample, and then potentially discovered exons are run through it for each smaple
    - the model uses the bp level count info at the boundary of an exon getting longer/shorter.
    - more often an exon is found th emore likely its actually there
- after I get this working, then ill add the downstream stuff back later

02/12/19 Changes
-all software versioned in config file
-added hmmscan protein domain search

-01/22/19
    - use gffcompare gtf not stringtie-merge gtf because gffcompare is significantly better than stringtie at mapping
    back to genes. GFFcompare found 20K novel tx vs 18K on st-merge, with the same number of transcript.
    - at the initial merge step with stringtie, filteing out transcripts with at least 1 tpm in a third of the samples

01/15/19 Changes
- converted STAR rules into shell from python
- made outputs more organized
- created a synthetic set by sampling 5 samples from all tissues >10 samples, incliding eye tissues, might wanna change that;
    using synthetic set as body set for gtf and rmats
- making mulitple tissue comparisons against the synth for rMATs instead of pair wise comparisons
- restructured outputs so things are a little more organized
- only using paired samples for rmats
- finally got rid of the shitty salmon command
-Reminder that STAR makes the bam even if the alignment fails
-following CHESS paper - run each bam individually through stringtie, then merge to a tissue level, then merge into 1
 use gffcompare at each meerge step;

-12/14/18
    -st-merge at at it default tpm cut off did nothing, so now going to do what chess ppl did and filter it at 1tpm per tissue

-12/13/18
    - tried GFFcompare on all samples first gave 300k tx's but salmon couldn't map to them, so will now use
      stringtie merge on a per tissue level, which will cut out a lot of transcripts, then merge with gffcompare.
    - moved gffread > tx into its own rule


**************REMEMBER TO CHANGE THE WORKING DIR IN THE CONFIG FILE IF YOU RERUN*********
'''
import yaml
def readSampleFile(samplefile):
    # returns a dictionary of dictionaries where first dict key is sample id and second dict key are sample  properties
    res={}
    with open(samplefile) as file:
        file.readline()#skip the first line because it has a header
        for line in file:
            info=line.strip('\n').split('\t')
            res[info[0]]={'paired':True if info[1]=='y' else False, 'tissue':info[2],'subtissue':info[3]}
    return(res)

def subtissue_to_gtf(subtissue, sample_dict):
    res=[]
    for sample in sample_dict.keys():
        if sample_dict[sample]['subtissue']==subtissue :
            res.append(f'st_out/{sample}.gtf')
    return (res)

def salmon_input(id,sample_dict,fql):
    paired=sample_dict[id]['paired']
    id= fql + 'fastq_files/' + id
    if paired:
        return('-1 {s}_1.fastq.gz -2 {s}_2.fastq.gz'.format(s=id))
    else:
        return('-r {}.fastq.gz'.format(id))

def subtissue_to_sample(subtissue, sample_dict):
    res=[]
    [res.append(f'data/salmon_quant/{subtissue}/{sample}/quant.sf') for sample in sample_dict.keys() if sample_dict[sample]['subtissue'] == subtissue ]
    return(res)

def make_tx_fasta_input(st):
    if st == 'all_tissues.combined':
        return 'data/gtfs/all_tissues.combined.gtf'
    elif st == 'pan_eye':
        return 'data/gtfs/pan_eye.gtf'
    elif st == 'gencode':
        return 'ref/gencode_comp_ano.gtf'
    else:
        return f'data/gtfs/final_tissue_gtfs/{st}.gtf'
def calculate_cov_input(sample_dict, eye_tissues):
    eye_tissues = set(eye_tissues)
    return [f'data/gene_cov/{id}_cov.per-base.bed.gz' for id in sample_dict.keys() if sample_dict[id]['subtissue'] in eye_tissues ]


with open(config['file_yaml']) as fyml:
    files=yaml.load(fyml,Loader=yaml.FullLoader)

#sample information
sample_file=config['sampleFile']
sample_dict=readSampleFile(config['sampleFile'])# sampleID:dict{path,paired,meta-data}
tissue_file=config['tissueFile']
subtissue_file=config['subtissueFile']
with open(tissue_file) as tf, open(subtissue_file) as sf:
    tissues= [line.strip('\n') for line in tf]
    subtissues= [line.strip('\n') for line in sf]
sample_names=sample_dict.keys()
#software version info
salmon_version=config['salmon_version']
stringtie_version=config['stringtie_version']
STAR_version=config['STAR_version']
R_version=config['R_version']
VEP_version=config['VEP_version']
TransDecoder_version=config['TransDecoder_version']
samtools_version=config['samtools_version']
gffcompare_version=config['gffcompare_version']
mosdepth_version=config['mosdepth_version']
bedtools_version=config['bedtools_version']
bedops_version = config['bedops_version']
clinvar_data=config['clinvar_vcf']
file_yaml = config['file_yaml']
crossmap_version = config['crossmap_version']

#python_env=config['python_env']
#commonly used files/paths
working_dir=config['working_dir']
STARindex='ref/STARindex'
ref_tx_fasta=files['ref_tx_fasta']
ref_GTF=files['ref_GTF']
ref_genome=files['ref_genome']
fql=config['fastq_path']
bam_path=config['bam_path']
stringtie_full_gtf='data/gtfs/all_tissues.combined.gtf'
hmmer_version=config['hmmer_version']
#win_size=config['window_size']
eye_tissues=['Retina_Adult.Tissue', 'Retina_Fetal.Tissue', 'RPE_Fetal.Tissue', 'RPE_Adult.Tissue', 'Cornea_Adult.Tissue', 'Cornea_Fetal.Tissue' ]

rule all:
    input:  
        'data/rdata/novel_exon_classification.Rdata',
        'data/rdata/all_tissue_quant.Rdata',  
        'data/novel_loci/hmmer/seq_hits.tsv', 
        'data/novel_loci/novel_loci_blast_results_nr.tsv', 
        'data/shiny_data/app_data/DNTX_db.sql',
        'data/rdata/pan_eye_quant.Rdata',
        expand('data/vep/{subtissue}/variant_summary.txt', subtissue = ['Retina_Fetal.Tissue', 'Retina_Adult.Tissue', 'gencode']), 
        calculate_cov_input(sample_dict, eye_tissues)
    #'data/rmats/all_tissues_psi.tsv', 
    #expand('data/gtfs/raw_tissue_gtfs/{subt}.combined.gtf', subt=subtissues)
    # input:stringtie_full_gtf,'data/exp_files/all_tissue_quant.tsv.gz','data/rmats/all_tissues_psi.tsv', 'data/rmats/all_tissues_incCounts.tsv', 'data/seqs/transdecoder_results/best_orfs.transdecoder.pep', 'data/rdata/novel_exon_classification.Rdata'

'''
****PART 1**** download files and align to genome
-still need to add missing fastq files
-gffread needs indexed fasta
-need to add versioning of tools to yaml{DONE}
04/08/2019 - added the mysql command to get a comprehensive refseq gtf from ucsc, which is somehow different than the one
from ncbi. see https://bioinformatics.stackexchange.com/questions/2548/hg38-gtf-file-with-refseq-annotations
'''

rule downloadAnnotation:
    output: 
        reftxfa=ref_tx_fasta,
        refgtf=ref_GTF, 
        ref_gen=ref_genome,
        prot_seq='ref/gencodeProtSeq.fa',
        gencode_gff='ref/gencodeGFF3.gff', 
        ensbl_gtf='ref/ensembl_ano.gtf', 
        refseq='ref/refseq_ncbi.gff3', 
        ucsc='ref/ucsc.gtf',
        clinvar_sum = 'ref/clinvar_variant_summary.txt.gz'
    shell:
        '''
        wget -O - {config[ref_tx_fasta_url]} | gunzip -c - > {output.reftxfa}
        wget -O - {config[refGTF_url]} | gunzip -c - > {output.refgtf}
        wget -O - {config[ref_genome_url]} | gunzip -c - > /tmp/gencodePA_tmp.fa
        wget -O - {config[refProtSeq_url]} | gunzip -c - > /tmp/gencodeProtSeq.fa
        wget -O - {config[refGFF3_url]q} | gunzip -c > {output.gencode_gff}
        wget -O - {config[ensembl_gtf_url]} | gunzip -c - > {output.ensbl_gtf}
        wget -O - {config[refseq_ncbi_url]} | gunzip -c - > {output.refseq}
        wget -O {input.clinvar_sum} {config[clinvar_variant_url]}
        module load {config[mysql_version]}
        module load {config[ucsc_version]}
        mysql --user=genome --host=genome-mysql.cse.ucsc.edu -A -N -e "select * from refGene" hg38 |\
        cut -f2- | genePredToGtf -source=hg38.refGene.ucsc file stdin {output.ucsc}
        module load python/3.6
        python3 scripts/filterFasta.py /tmp/gencodePA_tmp.fa ref/chroms_to_remove {output.ref_gen}
        python3 scripts/clean_fasta.py /tmp/gencodeProtSeq.fa {output.prot_seq}
        module load {samtools_version}
        samtools faidx ref/gencode_genome.fa
        awk '$3 == "transcript"' ref/gencode_comp_ano.gtf | cut -f1,7,4,5 > ref/gencode_comp_ano_trim.tsv
        
        wget -O ref/hg38_fair+new_CAGE_peaks_phase1and2.bed.gz https://fantom.gsc.riken.jp/5/datafiles/reprocessed/hg38_latest/extra/CAGE_peaks/hg38_fair+new_CAGE_peaks_phase1and2.bed.gz
        wget -O /tmp/polya.bed.gz 'https://polyasite.unibas.ch/download/atlas/2.0/GRCh38.96/atlas.clusters.2.0.GRCh38.96.bed.gz'
        python3 /data/swamyvs/ChromosomeMappings/convert_notation.py  -c /data/swamyvs/ChromosomeMappings/GRCh38_ensembl2gencode.txt -f /tmp/polya.bed.gz | bedtools sort -i stdin > ref/polyAtlas_gencodechrom_hg38_sorted.bed
        
        wget -O - ftp://ftp.ebi.ac.uk/pub/databases/gencode/Gencode_human/release_19/GRCh37.p13.genome.fa.gz | gunzip -c - > ref/hg19_gencode_genome.fa 

        wget -O ref/hg19ToHg38.over.chain.gz http://hgdownload.soe.ucsc.edu/goldenPath/hg19/liftOver/hg19ToHg38.over.chain.gz

        '''

rule liftover_intronic_variants:
    input:
        files['intron_variant_panel']
    output:
        hg19_vcf = 'data/intron_variant_analyis/retinal_dystrophy_nc_variant_panel_hg19.vcf',
        hg38_vcf = files['intron_variant_hg38_vcf']
    shell:
        '''
        python3 scripts/panel_variants_to_vcf.py  \
            --panel {input} \
            --hg19genome {files[hg19_genome]} \
            --outvcf {output.hg19_vcf}
        module load {crossmap_version}
        crossmap vcf /data/swamyvs/ocular_transcriptomes_pipeline/ref/hg19ToHg38.over.chain.gz \
            {output.hg19_vcf} \
            {ref_genome} \
            {output.hg38_vcf}
        '''
rule clean_phylop_and_snps:
    input: 
        snps=expand('ref/snps/bed_chr_{chrom}.bed.gz', chrom=list(range(23))[1:]), 
        pp='ref/phylop_20/hg38.phyloP20way.bw'
    output:
        snps='ref/snps/hg38.snps.all.sorted.bed.gz', 
        pp='ref/phylop_20/hg38.phyloP20way.sorted.bed.gz'
    shell:
        '''
        rm -rf snp_tmp
        mkdir snp_tmp
        module load {config[ucsc_version]}
        module load {bedops_version}
        moulde load {bedtools_version}
        bigWigToWig {input.pp} | wig2bed | bedtools sort -i - | gzip -c - > {output.pp}
        for i in ref/snps/bed_chr_*.bed.gz ; do zcat $i | tail -n+2   > snp_tmp/snps.bed 
        awk '$2 != $3{OFS="\t"; print $0}' snp_tmp/snps.bed  > snp_tmp/good_snps.bed 
        awk '$2 == $3 {OFS="\t";print $1, $2, $3+1, $4, $5, $6}' snp_tmp/snps.bed | cat - snp_tmp/good_snps.bed | sort-bed --max-mem 24G - | gzip -c - > ref/snps/hg38.snps.all.sorted.bed.gz
        rm -rf snp_tmp
        '''

# This is manily for rerunning on biowulf,
rule build_gffread:
    output:'gffread/gffread'
    shell:
        '''
        git clone https://github.com/gpertea/gclib
        git clone https://github.com/gpertea/gffread
        cd gffread
        make release
        '''

rule build_STARindex:
    input: ref_genome, ref_GTF
    output:directory(STARindex)
    shell:
        '''
        module load {STAR_version}
        mkdir -p ref/STARindex
        STAR --runThreadN 16 --runMode genomeGenerate --genomeDir {output[0]} --genomeFastaFiles {input[0]} --sjdbGTFfile {input[1]} --sjdbOverhang 100

        '''

rule run_STAR_alignment:
    input: fastqs=lambda wildcards: [fql+'fastq_files/{}_1.fastq.gz'.format(wildcards.id),fql+'fastq_files/{}_2.fastq.gz'.format(wildcards.id)] if sample_dict[wildcards.id]['paired'] else fql+'fastq_files/{}.fastq.gz'.format(wildcards.id),
        index=STARindex
    output:temp(bam_path+'STARbams/{id}/raw.Aligned.out.bam'), bam_path+'STARbams/{id}/raw.Log.final.out'
    shell:
        '''
        id={wildcards.id}
        mkdir -p STARbams/$id
        module load {STAR_version}
        STAR --runThreadN 8 --genomeDir {input.index} --outSAMstrandField intronMotif  --readFilesIn {input.fastqs} \
        --readFilesCommand gunzip -c --outFileNamePrefix STARbams/$id/raw. --outSAMtype BAM Unsorted
        '''

rule sort_bams:
    input:bam_path + 'STARbams/{id}/raw.Aligned.out.bam'
    output:bam_path + 'STARbams/{id}/Sorted.out.bam'
    shell:
        '''
        module load {samtools_version}
        samtools sort -o {output[0]} --threads 7 {input[0]}
        samtools index -b {output}
        '''


'''
****PART 2**** build Transcriptome, and process
'''
rule run_stringtie:
    input:bam_path + 'STARbams/{sample}/Sorted.out.bam'
    output:'st_out/{sample}.gtf'
    shell:
        '''
        module load {stringtie_version}
        stringtie {input[0]} -o {output[0]} -l {wildcards.sample} -p 8 -G {ref_GTF}
        '''

rule calculate_cov:
    input:bam_path+'STARbams/{id}/Sorted.out.bam'
    output: 'coverage_files/{id}/cov.per-base.bed.gz'
    shell:
        '''
        module load {mosdepth_version}
        sample={wildcards.id}
        mosdepth coverage_files/$sample/cov {input[0]}
        '''


rule merge_gtfs_to_tissue:
    input:
        gtfs = lambda wildcards: subtissue_to_gtf(wildcards.subtissue, sample_dict)
    params:
        outdir = lambda wildcards: f'data/gtfs/raw_tissue_gtfs/{wildcards.subtissue}', 
        st_quant_dir = 'st_out/'
    output:
        filtered_gtf = 'data/gtfs/raw_tissue_gtfs/{subtissue}.combined.filtered.gtf',
        conv_tab = 'data/gtfs/raw_tissue_gtfs/{subtissue}.convtab'
    shell:
        '''
        module load {gffcompare_version}
        gffcompare -r {ref_GTF} -T  -p {wildcards.subtissue} -o {params.outdir} {input.gtfs}
        module load {R_version}
        Rscript scripts/filter_gtfs_by_tissue.R \
            --workingDir {working_dir} \
            --sampleTable {sample_file} \
            --subtissue {wildcards.subtissue} \
            --gtfDir {params.outdir} \
            --stringtieQuant {params.st_quant_dir}
        '''

rule merge_all_gtfs:
    input: 
        gtfs = expand('data/gtfs/raw_tissue_gtfs/{subtissue}.combined.filtered.gtf', subtissue= subtissues)
    params: 
        gffc_prefix='all_tissues'
    output: 
        all_tis_gtf=files['base_all_tissue_gtf'], 
        master_conv_tab=files['all2tissue_convtab'],
        pan_eye_gtf = files['pan_eye_gtf']
    shell:
        '''
        module load {gffcompare_version}
        gffcompare -r {ref_GTF} -T -p DNTX -o data/gtfs/all_tissues {input.gtfs}
        module load {R_version}
        Rscript scripts/merge_tissue_gtfs.R  \
            --workingDir {working_dir} \
            --filesYaml {file_yaml}    
        
        '''
# Rscript scripts/merge_tissue_gtfs.R

'''
fix the tissue specific gtfs so they have the all_tissue novel transcript ids(DNTX); make salmon exp Rdata with the new tissue specific gtf

'''

rule make_tx_fasta:
    input: 
        tool = 'gffread/gffread', 
        gtf = lambda wildcards: make_tx_fasta_input(wildcards.subtissue)
    output: 
        'data/seqs/{subtissue}_tx.fa'
    shell:
        '''
        ./gffread/gffread -w {output} -g {ref_genome}  {input.gtf}
        '''


'''
pulled this from the trinnotate pipeline, makes a gff of the of using protein translations
'''
rule run_trans_decoder:
    input:
        'data/seqs/all_tissues.combined_tx.fa'
    output:
        'data/seqs/transdecoder_results/all_tissues.combined_transdecoderCDS.gff3',
        'data/seqs/transdecoder_results/transcripts.fasta.transdecoder.pep'
    shell:
        '''
        rm -rf TransDecoder
        git clone https://github.com/TransDecoder/TransDecoder.git
        cd TransDecoder
        module load {TransDecoder_version}
        mkdir -p ../data/seqs/transdecoder_results/
        ./util/gtf_genome_to_cdna_fasta.pl ../data/gtfs/all_tissues.combined.gtf ../ref/gencode_genome.fa > transcripts.fasta
        ./util/gtf_to_alignment_gff3.pl ../data/gtfs/all_tissues.combined.gtf > transcripts.gff3
        TransDecoder.LongOrfs -m 60 -t transcripts.fasta
        TransDecoder.Predict --single_best_only  -t transcripts.fasta
        ./util/cdna_alignment_orf_to_genome_orf.pl \
            transcripts.fasta.transdecoder.gff3 \
            transcripts.gff3 \
            transcripts.fasta > ../data/seqs/transdecoder_results/all_tissues.combined_transdecoderCDS.gff3
        mv transcripts.fasta.transdecoder.*  ../data/seqs/transdecoder_results/
        '''
#
rule clean_pep:
    input:
        'data/seqs/transdecoder_results/transcripts.fasta.transdecoder.pep'
    output:
        pep='data/seqs/transdecoder_results/best_orfs.transdecoder.pep', 
        meta_info='data/seqs/transdecoder_results/pep_fasta_meta_info.tsv'#, len_cor_tab='data/seqs/len_cor_tab.tsv'
    shell:
        '''
        python3 scripts/clean_pep.py {input} {output.pep} {output.meta_info}
        '''
                #python3 scripts/fix_prot_seqs.py /tmp/tmpvs.fasta  {output.pep} {output.len_cor_tab}

rule process_and_annotate_master_gtf:
    input: 
        gtf = files['base_all_tissue_gtf'],
        gff = 'data/seqs/transdecoder_results/all_tissues.combined_transdecoderCDS.gff3',
        full_pep = 'data/seqs/transdecoder_results/best_orfs.transdecoder.pep'
    params:
        agat_tmp_file = 'tmp/agat/gtf_startstop_added.gff',
        path_to_final_gtfs = 'data/gtfs/final_tissue_gtfs/',
        path_to_filt_gtfs = 'data/gtfs/raw_tissue_gtfs/'
    output: 
        full_gtf = files['anno_all_tissue_gtf'],
        tissue_gtfs = expand('data/gtfs/final_tissue_gtfs/{subtissue}.gtf', subtissue = subtissues),
        tissue_det_dfs = expand('data/gtfs/final_tissue_gtfs/{subtissue}.detdf', subtissue=subtissues),
        classfile=files['exon_class_rdata'],                
        novel_loci_pep=files['novel_loci_pep'],
        gencode_dummy = 'data/gtfs/final_tissue_gtfs/gencode.gtf',
        novel_loci_txids = files['novel_loci_txids'], 
        novel_loci_bed = files['novel_loci_bed']
    shell:
        '''
        agat_sp_add_start_and_stop.pl \
            --gff {input.gff} --fasta {ref_genome}  \
            --out  {params.agat_tmp_file} 
        
        module load {R_version}
        Rscript scripts/annotate_and_make_tissue_gtfs.R \
            --workingDir {working_dir} \
            --fileYaml {file_yaml} \
            --agatGff {params.agat_tmp_file}

        python3 scripts/select_entry_from_fasta.py \
            --infasta {input.full_pep} \
            --txToKeep {output.novel_loci_txids} \
            --outfasta {output.novel_loci_pep}
        
        cp {ref_GTF} {output.gencode_dummy}
        '''


rule run_vep:
    input:
        vcf =files['intron_variant_hg38_vcf'],
        gtf = 'data/gtfs/final_tissue_gtfs/{subtissue}.gtf'
    output: 
        gtf = 'data/gtfs/CDS_complete/{subtissue}.gtf.gz',
        variant_summary = 'data/vep/{subtissue}/variant_summary.txt'
    shell:
        '''
        rm -rf data/vep/{wildcards.subtissue}/*
        module load {samtools_version}
        module load {VEP_version}
        rm -rf {output.gtf} {output.gtf}.tbi 
        grep -v "#" {input.gtf} | sort -k1,1 -k4,4n -k5,5n -t$'\\t' | bgzip -c > {output.gtf}
        tabix -p gff {output.gtf}
        echo "running VEP"
        vep -i {input.vcf} --gtf {output.gtf} --fasta {ref_genome} -o {output.variant_summary}
        '''


rule get_genes_for_cov_analysis:
    input: full_gtf = files['anno_all_tissue_gtf']
    output: 'data/gene_cov/regions.bed'
    shell:
        '''
        for gene in ABCA4 IFT140 PROM1 RPGRIP1 
        do 
            grep $gene {input.full_gtf}|\
            cut -f1 -d';'  |\
            awk -v OFS='\t' '$3 == "exon"{print $1,$4,$5,$10}' |\
            tr -d '"'
        done > {output}
        '''

rule intersect_coverage:
    input:
        cov = 'coverage_files/{id}/cov.per-base.bed.gz', 
        bed =  'data/gene_cov/regions.bed'
    output:
        'data/gene_cov/{id}_cov.per-base.bed.gz'
    shell:
        '''
        module load {bedtools_version}
        bedtools intersect -a {input.cov} -b {input.bed} > {output}
        '''


rule build_salmon_index:
    input: 
        'data/seqs/{subtissue}_tx.fa'
    output: 
        directory('data/salmon_indices/{subtissue}')
    shell:
        '''
        module load {salmon_version}
        salmon index -t {input} -i {output} 
        '''

rule run_salmon:
    input: 
        fastqs=lambda wildcards: [fql+'fastq_files/{}_1.fastq.gz'.format(wildcards.sampleID),fql+'fastq_files/{}_2.fastq.gz'.format(wildcards.sampleID)] if sample_dict[wildcards.sampleID]['paired'] else fql+'fastq_files/{}.fastq.gz'.format(wildcards.sampleID),
        index='data/salmon_indices/{subtissue}'
    params: 
        cmd=lambda wildcards: salmon_input(wildcards.sampleID,sample_dict,fql),
        outdir=lambda wildcards: f'data/salmon_quant/{wildcards.subtissue}/{wildcards.sampleID}'
    output: 
        'data/salmon_quant/{subtissue}/{sampleID}/quant.sf'
    shell:
        '''
        id={wildcards.sampleID}
        module load {salmon_version}
        salmon quant -p 8 -i {input.index} -l A --gcBias --seqBias  --validateMappings {params.cmd} -o {params.outdir}
        '''

def all_quant_input(eye_tissues, sample_dict):
    res = []
    eye_tissues=set(eye_tissues)
    for key in sample_dict.keys():
        st = sample_dict[key]['subtissue']
        res.append(f'data/salmon_quant/{st}/{key}/quant.sf')
        res.append(f'data/salmon_quant/gencode/{key}/quant.sf')
        if sample_dict[key]['subtissue'] in eye_tissues:
            res.append(f'data/salmon_quant/pan_eye/{key}/quant.sf')
    return res 


rule merge_all_salmon_quant:
    input:
        all_quant_input(eye_tissues, sample_dict)
    params: 
        quant_path='data/salmon_quant/',
        out_dir = 'data/rdata/'
    output: 
        all_quant = files['all_tissue_quant'], 
        eye_quant = files['eye_quant'], 
        gencode_quant = files['gencode_quant']
    shell:
        '''
        module load {R_version}
        Rscript scripts/merge_salmon_quant.R  \
        --workingDir {working_dir} \
        --pathToQuant {params.quant_path} \
        --sampleTable {sample_file} \
        --outDir {params.out_dir}
        '''

'''

have to manually download this from UCSC
table browser > group=repeats, track=RepeatMasker
'''


rule blastp_novel_loci:
    input: 
        pep='data/novel_loci/novel_loci.pep'
    output: 
        swissprot_results='data/novel_loci/novel_loci_blast_results_swissprot.tsv',
        nr_results = 'data/novel_loci/novel_loci_blast_results_nr.tsv'
    shell:
        '''
        module load blast
        blastp -query {input.pep} -db /fdb/blastdb/swissprot  -max_target_seqs 250 -max_hsps 3 -outfmt "6 qseqid sseqid qlen slen nident evalue bitscore"  -num_threads 8 > {output.swissprot_results}
        blastp -query {input.pep} -db /fdb/blastdb/nr  -max_target_seqs 250 -max_hsps 3 -outfmt "6 qseqid sseqid qlen slen nident evalue bitscore"  -num_threads 8 > {output.nr_results}
        '''

rule build_pfm_hmmDB:
     params: url=config['pfam_db']
     output:'ref/hmmer/Pfam-A.hmm'
     shell:
         '''
         wget -O - {params.url} | gunzip -c - > {output}
         module load {hmmer_version}
         hmmpress {output}
         '''


rule run_hmmscan:
     input: 
        pfam='ref/hmmer/Pfam-A.hmm',  pep='data/novel_loci/novel_loci.pep'
     output:
        tab='data/novel_loci/hmmer/seq_hits.tsv',
        dom='data/novel_loci/hmmer/domain_hits.tsv',
        pfm='data/novel_loci/hmmer/pfam_hits.tsv'
     shell:
         '''
         module load {hmmer_version}
         hmmscan --cpu 24 --tblout {output.tab} --domtblout {output.dom} --pfamtblout {output.pfm} {input.pfam} {input.pep}
         '''


rule prep_shiny_data:
    input: 
        ano_gtf = files['anno_all_tissue_gtf'], \
        det_dfs = expand('data/gtfs/final_tissue_gtfs/{subtissue}.detdf', subtissue=subtissues), \
        tc2m = 'data/gtfs/all_tissues.convtab', \
        all_exp_file='data/rdata/all_tissue_quant.Rdata', \
        snps='ref/snps/hg38.snps.all.sorted.bed.gz', \
        pp='ref/phylop_20/hg38.phyloP20way.sorted.bed.gz', \
        gff3='data/seqs/transdecoder_results/all_tissues.combined_transdecoderCDS.gff3'
    params: 
        dd_stem='data/gtfs/final_tissue_gtfs/REPLACE.detdf',
    output:
        db_file='data/shiny_data/app_data/DNTX_db.sql'  , 
        rdata_file='data/shiny_data/app_data/shiny_data.Rdata', 
        dl_data_dir= directory('data/shiny_data/dl_data/')
    shell:
        '''

        rm -rf data/shiny_data/debug
        rm -rf {output.dl_data_dir}
        mkdir -p {output.dl_data_dir}
        mkdir -p data/shiny_data/debug
        touch data/shiny_data/debug/no_cds_bu_marked_as_pc.txt
        touch data/shiny_data/debug/multi_tx_under_same_id.txt
        touch data/shiny_data/debug/failed_to_find_CDS_start_or_end.txt
        touch data/shiny_data/debug/special_is_funky.txt
        touch data/shiny_data/debug/bad_genes.txt
        touch data/shiny_data/debug/igap_below_gap.txt
        module load {R_version}
        module load {bedtools_version}
        Rscript scripts/prep_data_for_shiny.R \
            --workingDir {working_dir} \
            --ddStem {params.dd_stem} \
            --filesYaml {file_yaml}
        for gtf in data/shiny_data/dl_data/*.gtf 
        do 
            stem=${{gtf::-4}}
            fasta=${{stem}}.fa
            ./gffread/gffread -w $fasta -g {ref_genome} $gtf
            zip ${{stem}}.zip $gtf $fasta
        done 
        '''

#note: need to to copy gtf instead of symlinking because snakemake wigs out about times stamps 
